# Network Inference from Multi-omic Data Uncovers Dynamic Transcriptional Regulation Modules in Pathogenic Fungus *Fusarium graminearum*

**DOI:** 10.1101/858498

**Authors:** Li Guo, Mengjie Ji, Kai Ye

## Abstract

The filamentous fungus *Fusarium graminearum* causes devastating crop disease and produces harmful mycotoxins worldwide. Understanding the complex *F. graminearum* transcriptional regulatory networks (TRNs) is vital for effective disease management. Reconstructing *F. graminearum* dynamic TRNs, an NP-hard problem, remains unsolved using commonly adopted reductionist or co-expression based approaches. Multi-omic data such as fungal genomic, transcriptomic data and phenomic data are vital to but so far have been largely isolated and untapped for unraveling phenotype-specific TRNs. Here for the first time, we harnessed these resources to infer global TRNs for *F. graminearum* using a Bayesian network based algorithm, “module networks”. The inferred TRNs contain 49 regulatory modules that show condition-specific gene regulation. Through a robust validation based on prior biological knowledge including functional annotations and TF binding site enrichment, our network prediction displayed high accuracy and concordance with existing knowledge, highlighted by its accurate capture of the well-known trichothecene gene cluster. In addition, we developed a new computational method to calculate the associations between modules and phenotypes, and discovered subnetworks responsible for fungal virulence, sexual reproduction and mycotoxin production. Finally, we found a clear compartmentalization of TRN modules in core and lineage-specific genomic regions in *F. graminearum*, reflecting the evolution of the TRNs in fungal speciation. This system-level reconstruction of filamentous fungal TRNs provides novel insights into the intricate networks of gene regulation that underlie key processes in *F. graminearum* pathobiology and offers promise for the development of improved disease control strategies.

## INTRODUCTION

Agricultural plants worldwide commonly suffer from devastating diseases caused by pathogenic fungi (Agrios, 2005), threatening food safety and human survival amid increasing global climate change. Fusarium head blight (FHB) caused by *Fusarium graminearum* (*Fg*) is a serious disease of cereal crops, reducing yield and polluting the grains with mycotoxins such as deoxynivalenol (DON) and zearalenone (ZEA) (Leslie *et al*., 2006). FHB pathogenesis is tightly controlled by host and pathogen gene regulatory networks (GRNs). For example, genes involved in *Fg* growth, infection and secondary metabolism are subject to fine regulation (Ma *et al*., 2013). Numerous studies have demonstrated that the expression of *Fg* genes related to pathogenesis, such as those encoding effectors (Brown *et al*., 2017) and cell wall-degrading enzymes (Carapito *et al*., 2008), is induced *in planta* but suppressed *in vitro*. Similarly, host genes involved in defense and immune response are induced during pathogen invasion (Pan *et al*., 2018). Understanding GRNs is fundamental in solving medical and agricultural problems (Emmert-Streib *et al*., 2014) caused by microbial infections. GRNs can inform disease control approaches by permitting the specific targeting of key pathogen regulators, as reported in recent studies (Koch *et al*., 2016). However, GRNs involved in FHB and mycotoxin production remain poorly understood.

Genes and gene regulators such as signaling proteins and transcription factors (TFs) are interconnected in GRNs. Many studies have attempted to dissect GRNs using a reductionist approach by analyzing the gene expression profiles of *Fg* mutants (Lysoe *et al*., 2011; Son *et al*., 2016; Kong *et al*., 2018). Though conceptually valid, this approach is time-consuming and unrealistic as a method of decoding the highly complex GRNs of eukaryotic cells. Alternatively, protein interaction networks have been inferred using protein domain homology. For instance, Zhao *et al*. constructed *Fg* protein-protein interaction (FPPI) networks using protein domains that are conserved in *Fg* and *Saccharomyces cerevisiae* (*Sc*) (Zhao *et al*., 2009). Despite its usefulness in finding potentially interacting proteins, this approach infers a network based solely on protein sequence features and therefore lacks functional support. A more feasible approach is to use genome-wide expression data to deduce regulatory networks. For example, Kim *et al*. predicted gene co-expression networks involved in virulence of *F. verticillioides* using RNA-Seq data from the *FSR1* mutant (Kim *et al*., 2018). In addition, Liu *et al*. constructed a co-expression network based on gene expression data and the FPPI database, identifying several hub pathogenicity genes and subnetworks (Liu *et al*., 2010). These approaches have indeed produced valuable insights into *Fg* co-expression gene modules. However, co-expression does not necessarily indicate a true regulatory relationship. Recently, Lysenko et al. used gene expression data combined with data on protein interactions and sequence similarity to study the networks important for virulence (Lysenko *et al*., 2013). While integrating multiple sources of evidence is an improvement, it still relies on co-expression evidence and does not prove actual regulatory relationships. Furthermore, because the study focused on small gene sets that have an impact on virulence, a systemic view of regulatory networks is lacking.

Regulatory relationships are typically inferred from large genome datasets using computational methods built on mathematical models (Kim *et al*., 2009). Boolean networks, Bayesian networks, and Mutual Information have already proven to be powerful models for inferring regulatory networks (Friedman *et al*., 2000; Shmulevich *et al*., 2002; Zhao *et al*., 2016). Bayesian networks are probabilistic models that are ideal for studying regulatory relationships using noisy data such as gene expression profiles (Friedman *et al*., 2000). Therefore, these models have been frequently adopted in various GRN-inference algorithms. Previously, from a large collection of transcriptomic data, we reconstructed a global GRN for *Fg* using the machine-learning method MinReg (Pe’er *et al*., 2006) based on a Bayesian network model, successfully predicting 120 top regulators for 13,300 *Fg* genes (Guo *et al*., 2016). Despite the progress it represents, this first *Fg* GRN has obvious limitations. First, it mainly focuses on master regulators that control general rather than fungal-specific biological processes. Second, it is essentially static and offers little insight into how the networks adapt to various changes in endogenous and environmental stimuli. Without such knowledge, it is difficult to predict how gene regulation of diverse and specific biological processes operates and to find the *bona fide* regulators in the system.

TFs regulate gene transcription via binding to the promoter regions of target genes. Elucidation of transcription regulatory networks (TRNs) is a vital step in mapping global GRNs. Here, for the first time, we reconstructed global TRNs for *Fg* by applying a module network learning algorithm (Segal *et al*., 2003) to a large collection of transcriptomic data and integrating a phenomic database of *Fg* TFs that were reported previously (Son *et al*., 2011). The integration of phenomics and transcriptomics data in this study allows us to identify 49 module networks that are directly involved in the cellular processes that underlie key phenotypes in *Fg*, yielding the novel and crucial knowledge that “regulator X regulates target genes Y under condition Z”. Validation of the networks demonstrates the high accuracy of the inference. Association mining of the predicted module networks reveals links between gene modules and fungal phenotypes. The condition-specific TRNs significantly improve the resolution of the *Fg* transcriptional circuits controlling virulence, sexual reproduction and mycotoxin production, laying a vital foundation for the development of novel regimes to minimize FHB occurrence and mycotoxin contamination. The *Fg* module network (FuNet) is available for public query and downloading (https://xjtu-funet-source.github.io/FuNet/FuNet.html).

## MATERIALS AND METHODS

### Fungal transcriptomic and TF phenomic data

The *Fg* transcriptome data were downloaded from PLEXdb (www.plexdb.org) and from the Filamentous Fungal Gene Expression database (http://bioinfo.townsend.yale.edu/) (Table S1). The data and the normalization procedure have been described previously (Guo *et al*., 2016). The phenome data for *Fg* TFs were obtained from literature (Son *et al*., 2011) (Table S2). The expression data and a candidate regulator list of 170 TFs showing phenotypic changes in disruption mutants were used as the input data for module network inference.

### Module Network algorithm implementation

Modularized TRNs were inferred using a probabilistic method called “module network” (Segal *et. al* 2003). Using the input data, the algorithm determined both the partitioning of genes to modules and the regulation program for each module in an iterative manner. For each iteration, the procedure searched for a regulation program for each module and then reassigned each gene to the module whose program best predicted its behavior. These two steps were iterated until convergence was reached using the expectation maximization (EM) algorithm, thereby returning the predicted regulatory modules containing a set of regulators and target genes. Each module was represented as a decision tree that specified the conditions under which target genes were regulated by a particular regulator and whether the regulation was positive or negative (Segal *et al*., 2003). The search process was repeated three times, and the same modules and regulation programs were returned.

### Validation of module networks

The predicted modules were first validated based on the consistency between regulator phenotypes and target gene expression. Based on three major phenotypes of *Fg*, each module was evaluated for consistency between the regulator phenotype and the experimental conditions using the following three validation points: 1) sexual reproduction; 2) virulence; and 3) mycotoxin production. To quantify the consistency of the evaluation, we developed a scoring function (named *Score_vp_*) for each validation point; this function was defined as

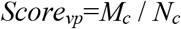

*N_c_* represents the total number of conditions included in this study, and *Mc* represents the number of conditions under which the regulator phenotype was matched with a corresponding condition directly related to the phenotype.

The regulatory modules were also validated based on conservation of *Fg* and *S. cerevisiae* TF binding site (TFBS). First, the 500-bp sequence upstream of the *Fg* genes of each module was extracted, and the *MEME* algorithm (Bailey *et al*., 2009) was used to search for conserved sequence motifs. The top five enriched motifs (ranked by E-value) were considered the candidate TFBS of each regulatory module. Each enriched TFBS was then compared to the *YEASTRACT* database of *S. cerevisiae* using *Tomtom* (Bailey *et al*., 2009) to find the conserved TFBS in budding yeast. The top conserved yeast motif for each *Fg* TFBS was then selected. With the existing knowledge of yeast TF-TFBS associations, the conserved yeast motifs identified through *Tomtom* identified corresponding TFs, which were denoted as “motif-deduced TFs” (MTFs). Second, to examine how many of these TFBS are potentially recognized by conserved TFs in *Fg* and *S. cerevisiae*, a BLASTp search was conducted in which the *Fg* regulators in each module were searched against the *S. cerevisiae* genome to find regulator orthologs (E-value < 1e-5); these were defined as “orthologous TFs” (OTFs). The OTFs were overlapped with MTFs to find conserved TFs that also potentially bind to conserved TFBS enriched in *Fg* regulatory modules. Based on the two separate analyses, a score was assigned to quantify the motif validation performance of each module:

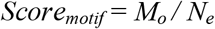

*M_o_* represents the number of motifs that showed conservation based on the above analysis, and *N_e_* represents the total number of enriched motifs per module.

Lastly, we validated each module by finding consistency between the functional annotations enriched in the module genes and the annotations associated with the enriched TFBS (deduced from *YEASTRACT*). First, GO enrichment was performed on the target genes from each module using MIPS Funcat (Ruepp *et al*., 2004) with the *Fg* genome as a reference. We then assigned functional annotations to each module by selecting the most highly enriched GO terms for each module. The annotations of the conserved enriched TFBS from each module were retrieved from *YEASTRACT* and compared to the GO terms enriched in the module genes. Following this analysis, we scored the module annotations matched with motif annotations as either 0 (no match) or 1 (match). We combined the scores from the above five validation points to obtain a total score; based on this score, the module was categorized as a high-confidence (>0.6), moderate-confidence (0.4∼0.6) or low-confidence module (<0.4). Modules with fewer than two validation points were not validated.

### Calculation of the module-phenotype association index

We developed an in-house computational method to accurately quantify the association between modules and phenotypes. We calculated a score called the association index (AI) for each module-phenotype association using multiple variables. The first variable (W*_ir_*) was the weight of the regulators derived from the number of conditions affected by the regulators specified by the regulation tree (Table S3). The number of experiments affected by each regulator in each module was used to obtain W*_ir_* (the weight of each regulator in each module; *i* =2, 3, 4…49 for modules M02, M03, M04…M49 and *r* =1, 2, 3…R for regulator 1, regulator 2, regulator 3…regulator R) according to the ratio of the number of experiments affected by the regulator. N*_ir_* indicates the number of conditions affected by the *r-th* regulator in the *i-th* module. W*_ir_* (0∼1) was calculated as follows:

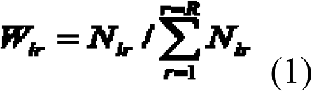

We then computed AI for each module-phenotype combination. Using the variable X*_irj_*, which could take a value of 0, 1 or −1, we could represent the influence of any regulator on a specific phenotype; the values 0, 1 and −1 indicated that the corresponding *r-th* regulator in the *i-th* module has no effect on, enhances or reduces, respectively, the *j-th* phenotype (Table S4). We calculated the AI (P*_ij_*) by multiplying the influence of regulator *r* on a phenotype *j*, denoted as X*_irj_*, by the weight of the corresponding regulator W*_ir_* and finally summing the product of all the regulators (*R*) in the module.

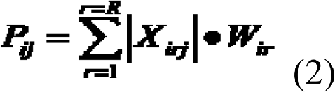

### Association mining

We created a correlation matrix of all modules based on phenotype associations. First, we filtered out minor associations (AI < 0.3) to capture major module-phenotype associations. The Pearson correlation coefficient (PCC) of each pair of modules was calculated and used to create a PCC matrix using the association indexes of the modules across phenotypes. Hierarchical clustering was used to find module clusters that are likely to contribute to similar phenotypes. Each cluster was subjected to detailed downstream examination.

### Network compartmentalization analysis

We identified 9,700 orthologous genes as core genes conserved among the three *Fusarium* sister species *Fg, F. verticillioides* and *F. oxysporum* (Ma *et al*., 2010). In total, 3,600 *Fg* genes lacking orthologous sequences in the sister species were loosely defined as *Fg* lineage-specific (LS) genes. We compared the observed ratio of LS and core genes in each predicted regulatory module to the expected ratio for the *Fg* genome using two-tailed Fisher’s exact test to determine whether there is an enrichment of LS or core genes. A threshold p-value < 0.05 was applied to determine whether the module was enriched (either LS or core) or not (mix).

### Network visualization

Cytoscape (version 3.6.1) (Shannon *et al*., 2003) and Gephi (version 0.9.2) (Bastian *et al*., 2009) were used for network visualizations. For building weight-based networks, modules and regulators are presented as nodes, and the weight of the regulator (W*_ir_*) in each module was used as the connection value for the edges. For phenotype-module networks, association indexes were used as the connection value of the edges.

### Network and code availability

The source codes used in this study are available at https://github.com/XJTU-FuNet-source/FuNet/tree/master/module-network_code-and-tool. The module networks of *Fg* are available at https://xjtu-funet-source.github.io/FuNet/FuNet.html.

## RESULTS

### 1. *Fusarium graminearum* module network inference

To infer condition-specific TRNs in *Fg*, we applied the module network algorithm (Segal *et al*., 2003) to a public dataset of *Fg* transcriptomic profiles spanning 67 different experimental conditions, including sexual reproduction and plant infection (Table S1). In addition, we used a set of candidate regulators consisting of 170 TFs that were previously functionally associated with key fungal phenotypes available in FgTFPD (*Fg* TF phenotype database) (Fig. 1) (Son *et al*., 2011). Combined expression data and phenotype-associated candidate regulators were used as input data for the module network algorithm to reconstruct TRNs in *Fg* (Fig. 1). Searching iteratively, the algorithm discovered 49 *Fg* gene regulatory modules, 48 of which had predicted regulators (Fig. 2a; Fig. S1; Table S5-S6).

**Figure 1.**
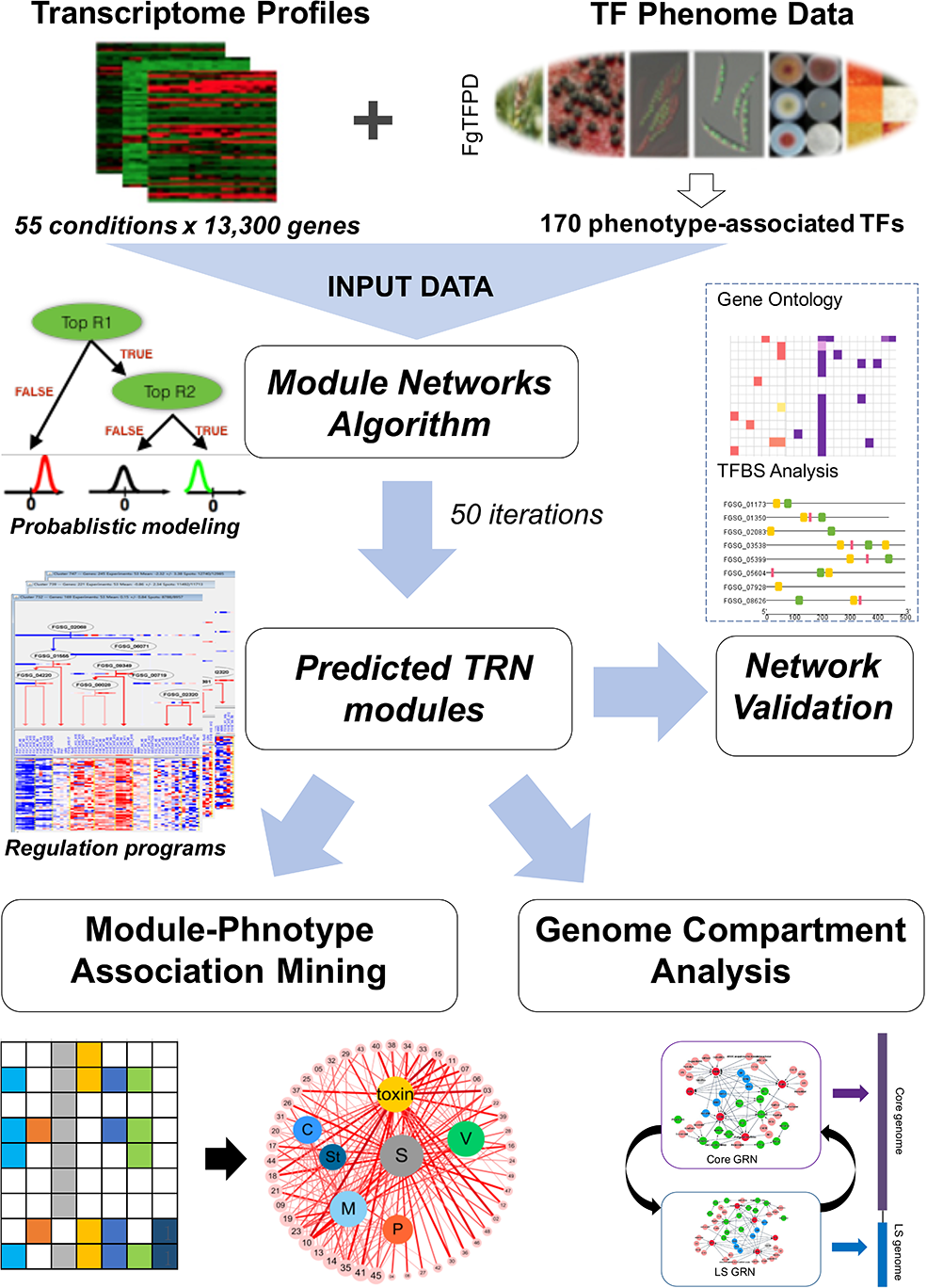
System framework and main steps in constructing a modularized gene regulatory network of *Fusarium graminearum*. Combined input consisting of gene expression data and information on candidate regulators was provided to the module networks algorithm (Segal *et al*. 2003). The inferred modules were then subjected to a robust validation process involving functional annotations and transcription factor binding site analysis. Then, the modules were analyzed for their phenotypic associations using an in-house computational method. Finally, the genome compartmentalization of the regulatory network was analyzed to provide evolutionary insights into the gene circuits.

**Figure 2.**
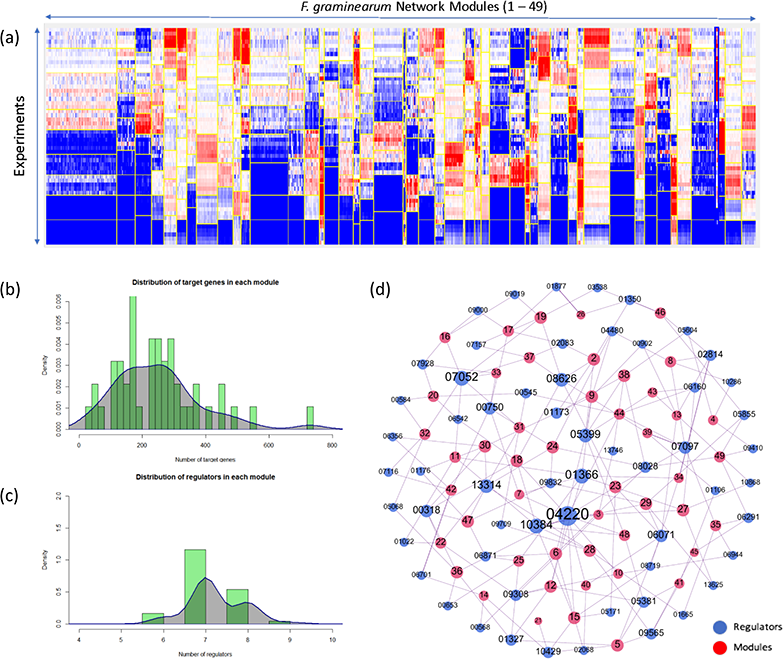
Overview of the module networks predicted for *Fusarium graminearum*. **(a).** Overview of inferred *F. graminearum* regulatory modules. The columns in the heatmap represent *Fg* genes, and the rows represent the experimental conditions in the gene expression data. Modules are delimited by vertical yellow lines. Red and blue represent gene activation and suppression, respectively. **(b).** Distribution of the number of target genes per module. Y axis represents the kernel density of target genes per module (X-axis). **(c).** Distribution of the number of regulators per module. Y axis represents the kernel density of regulators per module (X-axis). **(d).** An unweighted network of modules and regulators; the blue and red nodes represent regulators and modules, respectively. For clarity of visualization, the prefixes for regulator gene ID (“FGSG_”) and module (“M”) are omitted.

Each of the 48 modules is a regulatory program composed of various regulators, target genes and the expression profiles of target genes as a function of the expression level of the regulators. The regulatory program is presented in a decision-tree structure that defines the behavior of each regulator and the conditions under which the regulation takes place (Fig. S1). Overall, we predicted 117 regulators for 48 modules in *Fg*. The average numbers of target genes and regulators for each module are 268 and 7, respectively, with standard deviations of 212.43 and 1.24, respectively (Fig. 2b and 2c). The regulator-module association network (Fig. 2d) showed that 42 regulators were associated with only one module and that 75 regulators were associated with two or more modules. Five regulators including two ASPES proteins FGSG_04220 (13) and FGSG_10384 (7), a C2H protein FGSG_07052 (7), an HMG protein FGSG_01366 (8) and a Zn_2_C_6_ DNA binding protein FGSG_08626 (7) function as hub regulators associated with the greatest number of modules (Fig. 2d). Unsurprisingly, these regulators are highly pleiotropic, especially FGSG_04220 and FGSG_10384, whose deletion mutants are defective in the majority of phenotypes assayed (Table S2) (Son *et al*., 2011).

### 2. Validation using prior knowledge proves the high credibility of *Fg* module networks

Following the network inference, we assessed its reliability based on its consistency with prior knowledge. We scored each module by evaluating its performance based on multiple pieces of evidence, including regulator phenotypes, experimental conditions, gene annotations and cis-regulatory elements (Materials and Methods). A module was considered high-confidence, moderate-confidence or low-confidence depending on the validation score (Table S7). After discarding 16 modules for which there was little evidence, we identified 14 high-confidence, 13 moderate-confidence and 6 low-confidence modules. Overall, the high- and moderate-confidence modules account for 81.8% of the evaluable modules (Table S7), showing that our network inference has achieved a high degree of credibility. The following are examples of validation results that indicate the high credibility of our predicted modules.

#### 2.1. High concordance between regulator phenotypes and condition-specific gene regulation

Transcriptional regulators and their target genes are usually involved in the same biological processes. Based on this general premise, we validated all predicted modules by evaluating the concordance between the phenotypes associated with the top regulators in each module and our predicted condition-specific regulation using TF-phenotype associations and expression data associated with the experimental conditions. Since our predicted regulatory programs specify the regulators and the conditions under which the regulation occurs, the inferred relationship is accurate if a regulator associated with a phenotype and its target genes are both activated or suppressed under the experimental conditions that result in the phenotype. For simplicity and clarity, we focused our validation on the three largest groups of experimental conditions included in our data: sexual reproduction, plant infection and secondary metabolism (Table S1); these groups correspond to the sexual reproduction, virulence and mycotoxin (DON and ZEA) production phenotypes in FgTFPD, respectively. We found that in 45 of 48 modules with predicted regulators (Table S6), the top regulator has an effect on one or more of the phenotypes associated with sexual reproduction, virulence (plant infection) or mycotoxin production (secondary metabolism). In 34 of the 45 modules, the top regulator activates or suppresses the target genes under experimental conditions related to specific phenotypes (Table S7). Three specific examples, each concerning a phenotype, are provided below.

Firstly, in 76% of the modules whose top regulators are associated with sexual reproduction, regulation of the module genes by the top regulator was found under at least one sexual reproduction condition; for over 50% of the modules, the regulation occurred under half of the corresponding conditions (Table S7). For example, the top regulator of M30 (FGSG_06356) is essential for sexual reproduction. Our prediction showed that this TF and M30 genes were highly expressed under all sexual reproduction conditions. Secondly, in 57% of modules whose top regulators are associated with virulence, regulation of the module genes by the top regulator was found under at least one plant infection condition, and in nearly 30% of these modules, regulation occurred under half of the plant infection conditions (Table S7). For example, the top regulator of M16 (FGSG_07928) is essential for virulence, and our prediction showed that FGSG_07928 and the M16 genes were highly expressed under 62.5% of plant infection conditions. Thirdly, in 70% of the modules whose top regulators are associated with mycotoxin (DON or ZEA) production, regulation of module genes by the top regulator was found under 50% or more of the conditions that lead to mycotoxin induction (Table S7). For example, the top regulator for M46 (FGSG_03538) is essential for DON production, and our prediction showed that FGSG_03538 and the M46 genes were highly expressed under all mycotoxin induction conditions. In summary, 24 of the 32 predicted top regulators (75%) for 34 of the 48 predicted modules (70%) showed high concordance between the regulator phenotype and condition-specific gene regulation.

#### 2.2. Most predicted regulatory modules have functionally conserved TF binding sites

TFs regulate genes via binding to upstream cis-regulatory gene regions. Co-expressed genes (*e.g.*, genes in the same regulatory module) typically share TF binding sites (TFBS) that are recognized by one or more TFs. Therefore, we validated the predicted *Fg* (*Fg*) network modules by finding enriched TFBS in each module. Using the *MEME* algorithm, we first identified the top five enriched motifs (E-value < 0.05) within 500 bp upstream of the coding sequences for all *Fg* genes within a module (Table S7). Overall, 47 of 49 modules (96%) have significantly enriched motifs, and 34 (70%) have at least three enriched motifs (E-value <0.05). We then compared these significantly enriched motifs with the budding yeast *S. cerevisiae* (*Sc*) TFBS database *YEASTRACT* using *Tomtom* to functionally annotate these *Fg* motifs using conserved *Sc* motifs (Fig. 3; Table S8). To determine whether the enriched motifs were consistent with the biological functions of each module, we identified the most significantly enriched GO terms for each regulatory module (Fig. 3; Table S9) and compared the GO terms with the functional annotations of the significantly enriched *Fg* motifs. We found that the functional annotations of 27 of the 49 modules (55%) matched in the enriched TFBS and GO enrichment (Fig. 3; Table S10). For example, we found a functional match between M46 target genes and one of their enriched TFBS (*Yrm1p*) (Fig. 3); both are related to detoxification and multidrug resistance. This is consistent with the fact that M46 was highly associated with the mycotoxin production phenotype, as shown in later sections.

**Figure 3.**
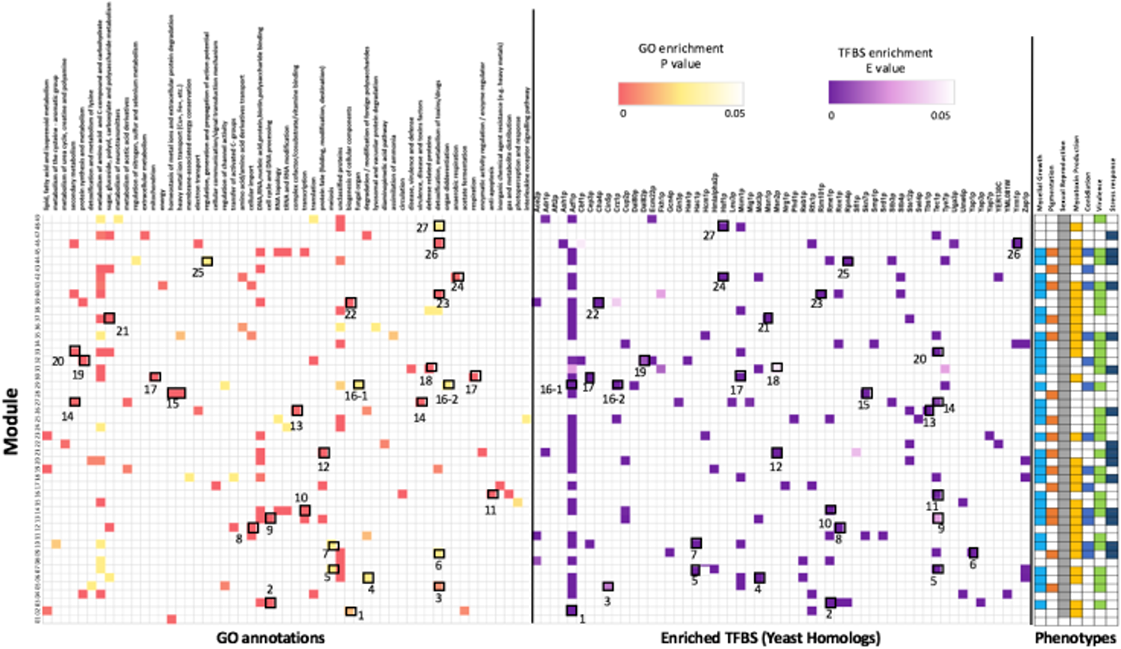
Gene Ontology (GO) annotation and transcriptional factor (TF) binding motifs enrichment in the predicted regulatory modules. GO enrichment was conducted for genes in each module, and major function associations were then assigned to each module using the most enriched GO terms. Each module is annotated with 55 main functional descriptions (P-value < 0.05). The *MEME* algorithm was used to find TF motif enrichment in the 500-bp region upstream of the *F. graminearum* genes sequence for each module (E value < 1e-5) followed by motif comparison with the budding yeast TF motif database using *Tomtom* (Materials and Methods). The heatmap on the left shows the significantly enriched GO terms per module; the color bar is scaled according to the P-value. The heatmap in the middle shows the motif enrichment results sorted into a total of 56 motifs; the color bar is scaled according to the E-value. The heatmap on the right shows the phenotype associations of each module based on association index analysis.

Secondly, to examine whether the functional conservation in TFBS was achieved through the conservation of regulator genes, a BLASTp search was conducted in which predicted *Fg* regulators were searched against the *Sc* genome to identify orthologous regulators (E-value < 1e-5). We then compared these yeast regulator orthologs with the *Sc* TFs derived from the *Sc* TFBS homologous to the enriched *Fg* TFBS. By overlapping the regulators identified in the two separate analyses, 10 different regulators regulating 36 modules (Table S11) were found, suggesting that not only did the predicted *Fg* regulators in 70% of the modules have conserved *Sc* homologs but also that conserved TFBS are likely associated with these fungal TF homologs. For example, motif enrichment shows that modules M06, M24, and M38 were enriched in a common TFBS (*Azf1p*) that is likely bound by YMR019W in *Sc*. One of the predicted regulators of module FGSG_08028 was orthologous to YMR019W (E-value = 1.46e-7). Another example is M39; this module is enriched in TFBS (*Ace2p*), which is likely bound by YLR131C in *Sc*. Interestingly, one of the predicted regulators of module FGSG_01341 is orthologous to YLR131C (E-value = 3.34e-17). The *YEASTRACT* database showed that this conserved TFBS and TF are involved in the biogenesis of cellular components, and this was captured by the GO enrichment for *Fg* module genes.

#### 2.3. Predicted regulatory modules captured the best-known TRN model in Fg

The best-understood model of a transcriptional regulation network in *Fg* is that of the trichothecene biosynthesis gene cluster, known as the Tri-cluster. Previous gene functional studies identified two TFs, *Tri6* (FGSG_03536) and *Tri10* (FGSG_03538), that reside within the cluster and regulate the expression of the entire cluster (Seong *et al*., 2009). Deletion of *Tri6* or *Tri10* in *Fg* abolished fungal production of trichothecene mycotoxins such as DON and its derivatives (Hohn *et al*., 1999; Tag *et al*., 2001). Remarkably, our network inference yielded a 44-gene module (M46) capturing the majority (9 of 12) of the Tri-cluster genes and accurately predicted *Tri6* and *Tri10* as the top two regulators of this module (Fig. 4a). To validate this module, we overlapped the module gene list (Table S5) with previously published lists of the genes that are down-regulated in *ΔTri6* and *ΔTri10* mutants (fold change ≥ 2) (Seong *et al*., 2009) as well as with lists of Tri6-binding genes identified by ChIP-Seq (Nasmith *et al*., 2011) and found that 34%, 22% and 11.3% of our module genes were included in the *ΔTri6* and *ΔTri10* down-regulated genes and *Tri6* ChIP-seq targets, respectively (Fig. 4b). Most of the overlapped genes were Tri-cluster genes. Furthermore, GO enrichment showed that this module was enriched in terms such as “secondary metabolism”, “detoxification involving cytochrome P450”, “sesquiterpenes metabolism”, “isoprenoid metabolism”, and others (Fig. 4c), consistent with the idea that this module regulates both primary and secondary metabolism as well as self-protection by *Tri10* and *Tri6* (Tag *et al*., 2001). Notably, our prediction specified that the target genes were activated by *Tri10* and *Tri6* during wheat and barley head infection, wheat coleoptile and crown rot infection, and toxin-inducing conditions but suppressed during sexual reproduction. In addition, M46 captured the full eight-gene cluster (FGSG_08077 ∼ FGSG_08084) for the biosynthesis of the *Fusarium* mycotoxin butenolide, which is toxic to animals (Liu *et al*., 2007). We predicted that this gene cluster is regulated by *Tri6* and *Tri10*, providing further evidence that the two TFs might be global regulators of secondary metabolism including trichothecene production, as previously reported (Seong *et al*., 2009; Nasmith *et al*., 2011).

**Figure 4.**
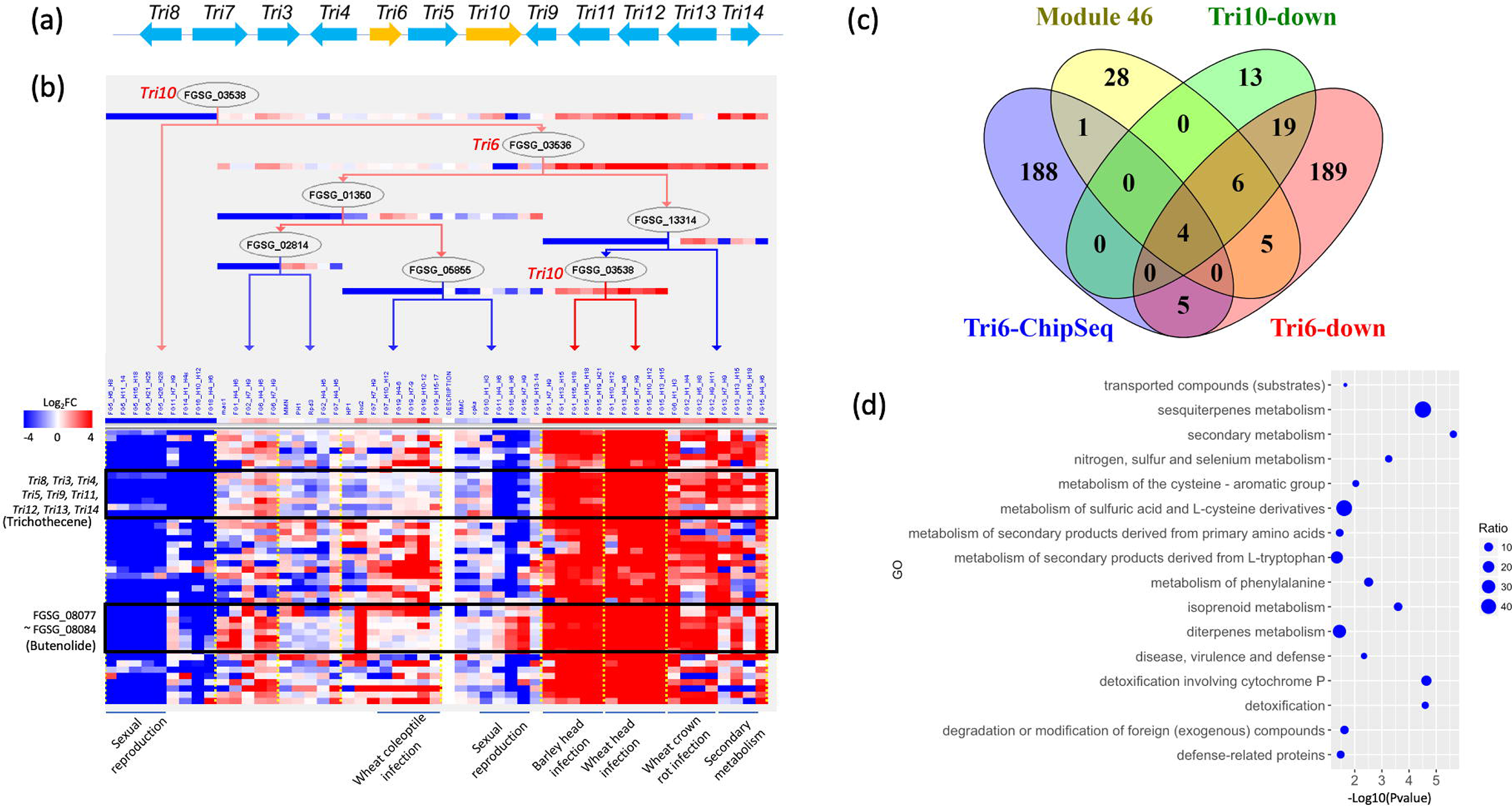
High accuracy of network inference highlighted by its capture of the trichothecene biosynthetic gene cluster. **(a).** Diagram of the trichothecene biosynthetic gene cluster (*Tri*-cluster) in *F. graminearum*. **(b).** Regulation program of the predicted module M46. The predicted regulators are arranged in a decision-tree structure that specifies the conditions under which the regulation occurs; these are delineated by the yellow vertical dotted lines. The heatmap underneath the tree shows the expression values of genes (rows) under different experimental conditions (columns). The experimental conditions are grouped into several major groups that are underlined; these are labeled at the bottom of the heatmap. The black empty box highlights all the *Tri*-cluster genes (labeled on the left) predicted in this module. **(c).** Venn diagram displaying the overlap of M46 genes with the previously published down-regulated genes of *ΔTri6* and *ΔTri10* mutants as well as *Tri6*-binding genes identified by ChIP-seq. **(d).** Dot plot showing the functional enrichment of M46 genes. The Y axis represents the significantly enriched GO terms. The X axis represents the P value (-Log10 transformed). The size of each dot is proportional to the gene ratio (Test/Reference), which was calculated as the number of genes in M46 (Test) divided by the number of genes in the *F. graminearum* genome (Reference) that are annotated with the GO term.

### 3. Computational method revealing module-phenotype association networks

To understand which modules are responsible for regulating key phenotypes in *Fg*, we developed a computational framework and used it to calculate an association index (AI) between each module and each phenotype (Fig. 5a; Table S12) using digitalized regulator phenotypes (Table S4) weighted by regulator hierarchical positions on the decision tree (Table S3) and the fraction of phenotype-matching conditions being regulated (Materials and Methods). We found that the module-phenotype association was quite complex in that the majority of modules were associated with multiple phenotypes. Under a threshold of AI > 0.3, we removed minor associations and clarified the primary associations between modules and different phenotypes and built module-phenotype association networks (Fig. 5b). We found that sexual reproduction (49), mycelial growth (33), mycotoxin production (31) and virulence (30) were associated with the largest numbers of modules.

**Figure 5.**
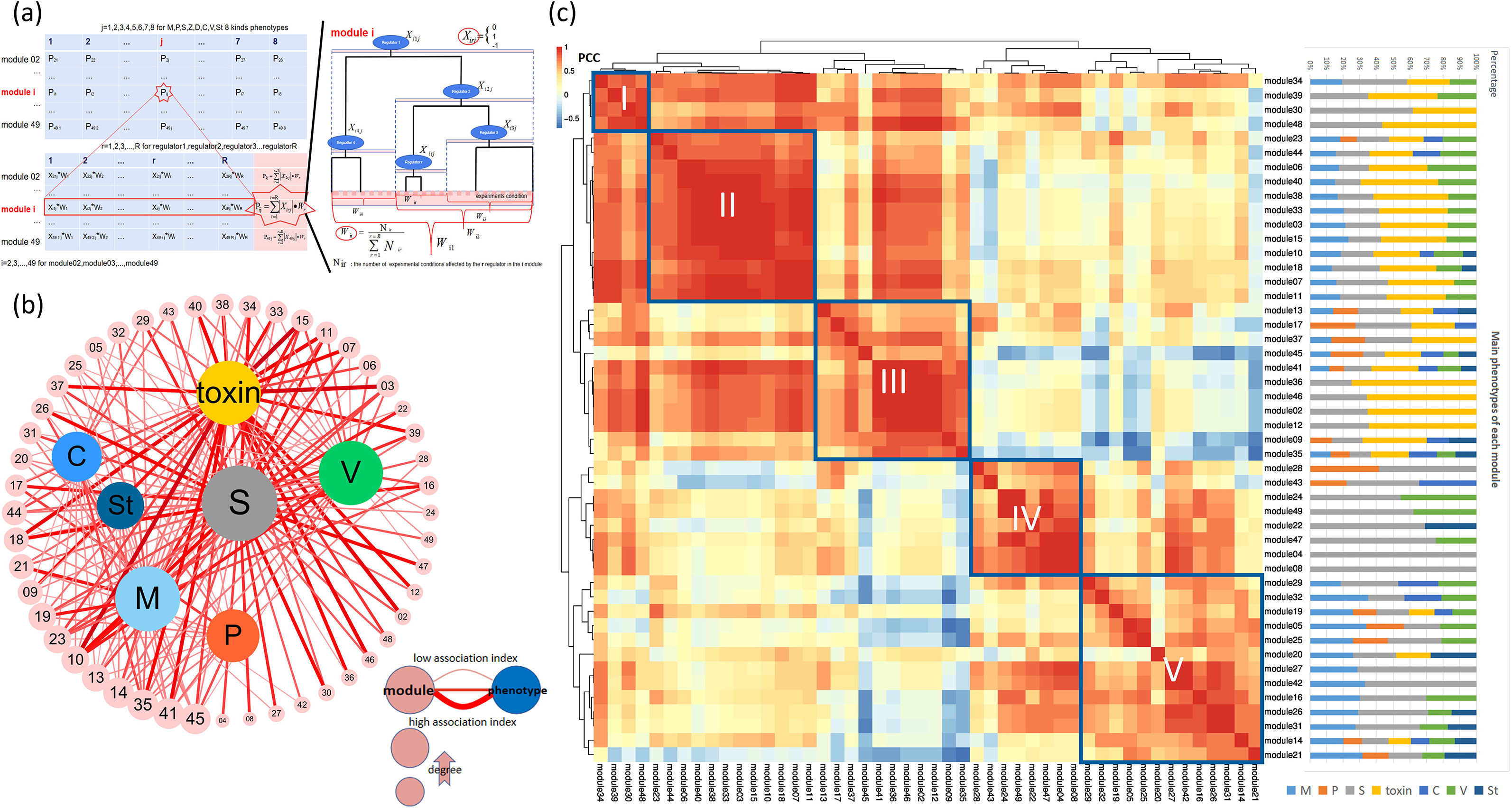
Computational analysis of module-phenotype associations. **(a).** Schematic demonstration of the computational method used to mine the module-phenotype association. The computational method calculates the association index based on the regulator’s phenotype, the regulator’s hierarchical position on the regulation tree, and the proportion of activated conditions in all experiments. **(b).** Module-phenotype association network based on the association index. The outer circle nodes represent modules, and the inner nodes represent phenotypes: M (mycelial growth), C (conidation), P (pigmentation), V (virulence), Toxin (DON and ZEA production), S (sexual reproduction) and St (stress response). The edges denote the associations between modules and phenotypes; the line sizes are proportional to AI of the association. **(c).** Correlation analysis of all predicted modules using Pearson correlation coefficients (PCC) based on the calculated association index. The modules are clustered using hierarchical clustering, and a heatmap of the PCCs (the scale bar denotes the PCC range) is then produced. The stacked bar graph adjacent to the heatmap summarizes the phenotype association within each module based on the association index.

Regulatory modules typically work as groups to control cellular functions. Therefore, we investigated the relationships among different modules with respect to their contribution to phenotypes. We measured the correlation among all modules by calculating the pairwise Pearson correlation coefficients (PCC) and then performed hierarchical clustering of the modules based on their PCCs. The analysis identified five major clusters of gene modules; each cluster showed a distinctive pattern of association with phenotypes (Fig. 5c). Cluster I included four modules (M34, M39, M30 and M48) in which sexual reproduction and mycotoxin production are major phenotypes as well as modules M34 and M39, which are mildly linked to mycelial growth and/or virulence. Cluster II contained 12 modules (M23, M44, M06, M40, M38, M33, M03, M15, M10, M18, M07 and M11) that are all linked to mycelial growth, sexual reproduction, mycotoxin production and virulence. Cluster III was composed of 11 modules (M13, M17, M37, M45, M41, M36, M46, M02, M12, M09, and M35) that are associated with pigmentation, in addition to other phenotypes. Overall, the phenotype associations of Clusters I to III were dominated by sexual reproduction and toxin production, and M30, M48, M36, M46, M12 and M02 were exclusively associated with these two phenotypes. Cluster IV contained eight modules (M28, M43, M24, M49, M22, M47, M04, and M08) whose phenotype associations were dominated by sexual reproduction and virulence and two modules that were exclusively associated with virulence. Cluster V contained 13 modules (M29, M32, M19, M05, M25, M20, M27, M42, M16, M26, M31, M14, and M21) associated with mycelial growth. However, except for the fact that M27 and M42 were mostly associated with mycelial growth and sexual reproduction, no phenotype associations were dominant.

### 4. TRN subnetworks that control key fungal phenotypes

From the association analysis results, we built subnetworks of *Fg* controlling virulence, sexual reproduction and mycotoxin production (Fig. 6). In these subnetworks, the nodes represent top regulators and modules, and the edges represent regulatory relationships weighted by the degree of the regulator’s influence on the module.

**Figure 6.**
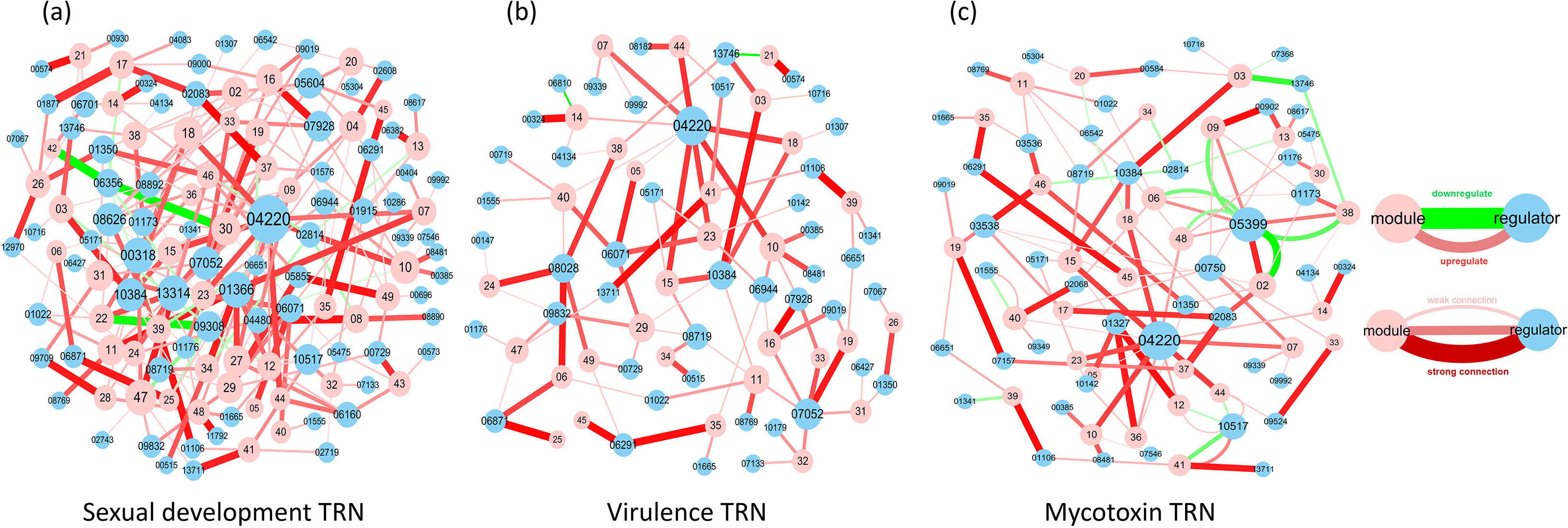
*Fusarium graminearum* subnetworks involved in the regulation of key phenotypes: sexual reproduction **(a)**, virulence **(b)**, mycotoxin production **(c)**. In each network, the edges represent the connections between the two nodes, which in turn represent the contributions of different regulators to the module. The red line represents a positive effect of the regulator on the corresponding phenotype, the green line represents a negative effect, and the thickness and color depth of the connection indicates the strength of the effect.

#### 4.1. Sexual reproduction TRNs

Sexual reproduction is critical for *Fg*, which produces ascospores as the primary inocula in the field. Based on the association index, we identified 10 modules (M30, M28, M24, M49, M22, M47, M04, M08, M27 and M42) that are highly associated with sexual reproduction and that have an association index of over 50% for all phenotype associations (Fig. 5c). The top regulators of the 10 modules were all essential for normal development of perithecia, and most act as positive regulators. For example, FGSG_04480 and FGSG_08890, the two top regulators of M08, activated target genes during sexual development (Fig. S1-8).

Interestingly, FGSG_08890 is a mating-type locus gene (*MAT-1-3*) and therefore potentially contributes to fungal mating by regulating sexual reproduction-related gene modules. In contrast, the two TFs FGSG_06356 and FGSG_09308 are negative regulators of sexual reproduction. For example, FGSG_06356, which is negatively associated with sexual development (Son *et al*., 2011), was predicted as a top regulator for M30 and M42. Consistently, our regulation program for M30 predicted that FGSG_06356 acts as a suppressor of M30 genes during sexual development (Fig. S1-30). Likewise, FGSG_09308 was predicted as a suppressor of M22 (Fig. S1-22), consistent with its previously reported phenotype. In addition, some regulators that had connections with many modules acted as hub regulators in sexual reproduction-associated networks, including FGSG_04220, FGSG_00318, FGSG_01366, FGSG_13314, FGSG_07052, FGSG_08626 and FGSG_10384 (Fig. 6a). Unsurprisingly, mutants in which these regulators are disrupted display defective sexual development (Table S2).

#### 4.2. Virulence TRNs

*Fg* is a pathogen of wheat and barley. To understand the transcriptional regulatory circuits in *Fg* infection, we built a network of gene modules and the top regulators associated with virulence using an AI threshold of > 0.3 (Fig. 6b). In total, 30 modules were associated with virulence (Fig. 5c); 5 of these (M06, M24, M49, M47 and M16) had the highest association with virulence, which contributed 25% to 45% of their phenotype associations. Four of the 30 modules, FGSG_04220, FGSG_07052, FGSG_08028 and FGSG_10384, were hub regulators of virulence. FGSG_08028, the top regulator for M06, M24 and M49, activated genes of the three modules during wheat plant infection. The previous report that a disruption mutant of FGSG_08028 loses pathogenicity but appears normal with respect to other phenotypes (Son *et al*., 2011) and our prediction that FGSG_08028 participates only in the virulence-associated TRN (Fig. 6b) suggest that this TF might exclusively regulate virulence. Notably, both positive and negative regulators were found in the virulence TRNs. For instance, FGSG_13314 and FGSG_00318, which are essential for pathogenicity, were the top two regulators positively regulating M47 genes under plant infection conditions (Fig. S1-47). Mutants of these TFs display abnormal development of perithecia and ascospores, consistent with their presence in the sexual reproduction TRN (Fig. 6a). On the other hand, the top two regulators of M16, FSGG_07928 and FGSG_09019, negatively regulate M16 gene expression during plant infection. However, disruption mutants of these TFs are nonpathogenic to wheat. GO enrichment of the M16 genes suggested that the module is highly enriched in anti-apoptosis processes. Loss of the function of its two main suppressors might contribute to decreased fungal apoptosis during infection, which is generally considered to be beneficial to pathogens during infection (Shlezinger *et al*., 2011). We further compared our predicted regulators with 70 existing genes reported in PHI-base whose mutants showed reduced or lost virulence (Urban M et al., 2017) to identify overlapped genes. Among these 70 virulence-associated genes reported in PHI-base, 63 were found in our total predicted regulators, 44 of which were included in virulence TRNs modules (Table S13), suggesting our network prediction has resulted in a virulence regulatory network that captured the majority (63%) of the known *Fg* virulence-associated genes in public domain.

#### 4.3. Mycotoxin TRNs

*Fg* produces toxic secondary metabolites, including DON and ZEA, that are harmful to livestock and to humans who consume the mycotoxin-containing maize products. The regulatory networks involved in fungal mycotoxin production are still poorly understood. Here, we found 31 TRN modules that were associated with mycotoxin production under the AI index threshold of 0.3 (Fig. 6c); six of these (M02, M12, M36, M46 and M48) were highly associated with mycotoxins in 55% ∼ 85% of all phenotypic associations (Fig. 5c). The majority of the Tri-cluster was captured by another mycotoxin-associated module, M46. Several regulators, including FGSG_04220, FGSG_05399, FGSG_00750, FGSG_10517, FGSG_01173, FGSG_03538 and FGSG_10384, acted as hub regulators in mycotoxin-associated modules. Not surprisingly, disruption mutants of these regulators show abnormal production of DON and ZEA (Table S2). Two of them, FGSG_05399 and FGSG_10157, are negative regulators of DON and ZEA, respectively. In addition, we predicted FGSG_05399 as the top negative regulator of M02 in the mycotoxin production and plant infection stages. Interestingly, GO enrichment suggested that the M02 genes are highly enriched for isoprenoid metabolism, which generates the precursors of trichothecenes. Although M02 did not include genes in the Tri-cluster, it contained a deacetylase gene, FGSG_03544, that is located near the Tri-cluster and is co-expressed with Tri-cluster genes (Sieber *et al*., 2014).

### 5. *Fg* module networks have core and lineage-specific (LS) compartments

A key question in evolution is how newly acquired genes or chromosomes integrate with existing or ancestral genomes in an organism and how the regulatory networks carried by two distinct compartments, i.e., the ancestral and the novel genetic materials, are accommodated to ensure the stability of the newly formed genome and its regulation during speciation. Previous comparative genomic analysis suggested that the *Fg* genome can be roughly divided into core and LS (Guo *et al*., 2016). To determine whether our predicted *Fg* module networks have a compartmentalized organization, we applied Fisher’s exact test to the genes in each module and identified modules significantly enriched for core and LS genomic regions. We found that eight modules (M06, M09, M25, M27, M34, M38, M46 and M48) showed significant enrichment of genes from LS regions, while 24 modules were enriched with genes from core regions (Fig. 7). Phenotypically, these eight LS modules were primarily associated with mycotoxin production, virulence and sexual reproduction (Fig. 6). Interestingly, among the 117 predicted regulators, only ten were located in LS regions. However, we found that four of them (FGSG_01327, FGSG_03536, FGSG_03538, and FGSG_08028) were exclusively associated with LS modules and that six were associated with other modules. Fisher’s exact test showed these ten LS regulators showed a significant bias of regulator-module association towards core or LS genomic regions (P-value < 0.05). FGSG_03536 (*Tri6*) and FGSG_03538 (*Tri10*) were the top two regulators of the LS module M46 (Fig. 4a), and FGSG_08028 was the top regulator of the LS modules M06 and M38. These four LS regulators, together with their associated LS modules, might be horizontally acquired and may have contributed to the speciation of *Fg* via specifically regulating the fungal biosynthesis of specific mycotoxins or plant infections. Lastly, a reciprocal regulation was found for regulators and modules across the LS and core genomes (Table S14), suggesting that there is communication between core and LS genomic regions and reflecting the evolutionary signature of regulatory programs involving both vertically and horizontally acquired gene regulatory elements.

**Figure 7.**
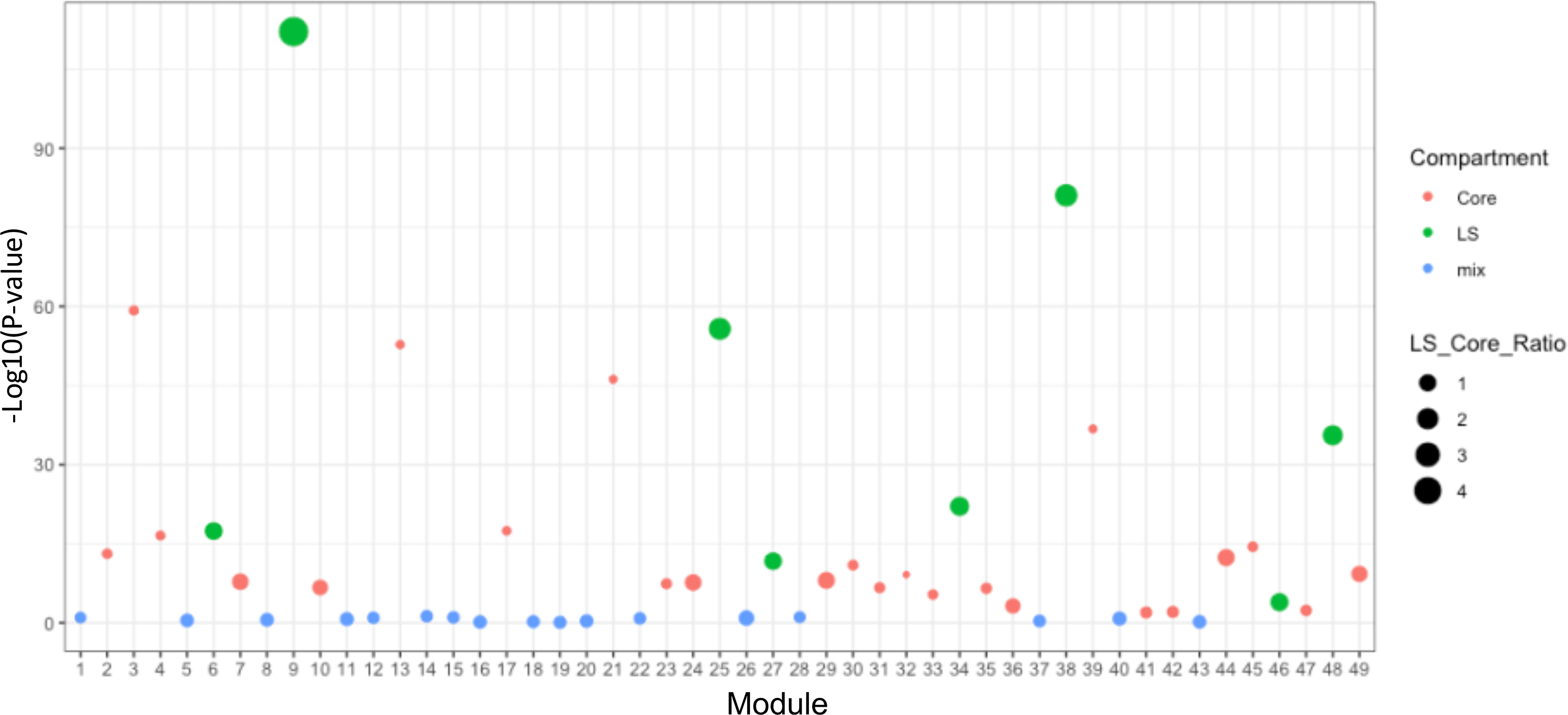
Compartmentalization of module networks into core and lineage-specific genome regions of *Fusarium graminearum*. Each of 49 *F. graminearum* modules was tested for enrichment of LS and core genes using two-tailed Fisher’s exact test. The dot plot summarizes the enrichment results; the X axis shows the module, the Y axis shows the test P-value for each module (-log10 transformed), and the solid horizontal line represents the threshold (P-value < 0.05). The color of the dot denotes the enriched compartment (green: LS; red: core; blue: not enriched) for each module. The size of the dot is proportional to the ratio of LS to core genes in each module.

## DISCUSSION

GRNs are the heart of every cellular function. Given its high complexity, the complete reconstruction of GRNs in multicellular organisms remains a daunting task. Traditional reductionist approaches and gene co-expression analysis have failed to solve this challenge in *Fg*. Here, we used a module network algorithm (Segal *et al*., 2003) to reconstruct *Fg* TRNs. The module network algorithm, which is a probabilistic method built on a Bayesian networks model, is ideal for dealing with noisy data such as gene expression profiles. Importantly, its strong performance in predicting dynamic and condition-specific gene regulation from large transcriptome datasets has been demonstrated (Segal *et al*., 2003). Furthermore, this TRN inference used phenotype-associated TFs as candidate regulators, allowing us to pinpoint the TRNs that directly mediate various fungal phenotypes. In contrast, previous GRNs were comprehensive but coarse, focusing on the master regulatory networks that control generic cellular functions. Lastly, we used phenomic data from TF knockout mutants as well as transcriptome data to discover phenotypically specific regulatory networks.

Given such technical advantages, the TRNs inferred here present two major improvements compared to previous GRNs. First, the TRNs predicted in this work improved the network resolution compared to our previous GRNs which inferred 120 regulators, each regulating 329 genes on average (Guo et al., 2016). In comparison, each module in the new TRNs contains an average of 268 genes and 7 regulators, significantly improving the network resolution. Previous GRNs predicted 44 transcription factors as regulators, while the TRNs yielded 117 TFs as regulators, suggesting the new TRNs found more regulators and therefore increased the resolution of regulatory networks involving TFs. Second, the TRNs are dynamic networks that specify the conditions under which a particular regulation occurs, unlike previous static GRNs. For example, we found that the TRNs of this work and previous GRNs shared eight regulators (Table S15). For these eight regulators, this TRN produced a dynamic network model that specifies the conditions under which a particular regulation occurs, while in previous static GRNs no such information existed. Third, we developed a new computational method for establishing associations between key fungal phenotypes and predicted network modules, while previous co-expression networks and GRNs lack any such association. The module-phenotype associations suggest that cellular phenotypes are most often controlled by complex networks of gene modules. Our association mining analysis also revealed major groups of modules that contribute primarily to specific fungal phenotypes, suggesting that biological processes in fungi are controlled by multiple intertwining gene modules. Finding the links between regulatory modules and phenotypes enriches our understanding of how fungal cells control their activities and, importantly, informs targeted approaches for suppressing specific gene modules and key regulators for disease control.

The TRN inference in this study significantly improves our understanding of transcriptional regulation in *Fg*. In previous functional studies, many TFs critical for fungal biology have been characterized. However, little is known about what target genes they regulate, which networks they use to regulate cellular functions, and, more importantly, when the regulation occurs and how it changes when the environment is altered. We identified 49 TRN modules in *Fg*. Each module represents a regulatory program in which a set of TFs regulates a number of target genes, and the hierarchical organization of these TFs reflects their influence on the module. Our biological validations suggested that our TRN inference achieved overall high performance and that it successfully associated TFs and their target genes. Importantly, our inference reveals the conditional specificity, the direction (positive vs. negative) and the strength of gene regulation, factors that are not addressed in previous *Fg* network studies.

*Fg* is a pathogenic fungus that is capable of producing harmful mycotoxins. Sexual reproduction plays a vital role in generating inocula for fungal infection. Therefore, understanding the regulatory networks that control virulence, sexual reproduction and mycotoxin production is a priority for research on *Fg* systems biology. For the first time, we identified TRN subnetworks in filamentous fungi that control three *Fg* phenotypes. Unlike studies that have focused on the contribution of a single gene or a set of co-expressed genes to phenotypes, we reconstructed TRN subnetworks that are strongly associated with each phenotype. The three subnetworks depicted the dynamic regulation of gene modules that are highly associated with each of the phenotypes. This work provides the best knowledge to date of the transcriptional regulation involved in these processes in *Fg* and lays an important foundation for further systems biology studies in the fungus. We previously reported that there appears to be a compartmentalization of *Fg* GRNs in core and LS genomic regions. The TRNs inferred here confirmed such compartmentalization but also provided novel insights into how the regulatory modules are organized into the core and LS compartments based on our identification of 8 and 24 modules that are significantly enriched in target genes from the LS and core genome regions, respectively. The top regulators in four of the eight LS-enriched modules are located in the LS genome, suggesting that the regulatory circuits associated with LS modules were very likely acquired with LS genome regions but that over time at least part of these acquired circuits were integrated into the ancestral circuits carried by the fungus.

Overall, we reconstructed dynamic TRNs from a large scale of *Fg* transcriptomic data using a module network algorithm. The overall accuracy of the network inference is high as demonstrated by validation using prior knowledge including functional annotation, TF binding site enrichment analyses and finally the deletion mutant transcriptome profiles in the well-established Tri-cluster regulatory model. Despite the progress reported here, this network remains an early step toward obtaining a complete understanding of authentic TRNs and eventually achieving its predictive power. As high-throughput sequencing technologies rapidly develop and costs continue to decrease, generation of additional large-scale genomic data such as RNA-seq and ChIP-seq will be needed to improve the network model and realize its predictive power in the future.

## ACKNOWLEDGMENTS

This project was supported by the National Natural Science Foundation of China (31701739, 31671372) and National Key R&D Program of China (Grant Nos. 2018YFC0910400 and 2017YFC0907500). L.G. is also supported by a China Postdoctoral Foundation Grant (2017M623188) and the Fundamental Research Fund of Xi’an Jiaotong University (1191329155).

## AUTHOR CONTRIBUTIONS

L.G. and K.Y. conceived and designed the project. L.G. and M.J. performed the network inference and validations. M.J. conducted the module-phenotype association analysis. L.G. and M.J. prepared the figures and tables. K.Y., L.G. and M.J. wrote the manuscript. L.G. and M.J. contributed equally to this work. All authors read and approved the manuscript.

## Supplementary Information

**Figure S1.** Regulatory programs for all 49 regulatory modules in *F. graminearum*.

**Table S1.** Gene expression data sets used to predict module networks of *F. graminearum* in this work.

**Table S2.** Phenotype summary of 170 transcription factors as candidate regulators extracted from the originally reported by Son *et al*. 2011 in PLoS Pathogens.

**Table S3.** Weight of each regulator (R) in each module.

**Table S4.** Digital representation of various regulator phenotypes.

**Table S5.** Lists of target genes in each module.

**Table S6.** Summary of 49 regulatory modules, including information on the number of target genes, the top regulators, and the top four enriched Gene Ontology terms (P value < 0.05).

**Table S7.** Summary of regulatory module validation results.

**Table S8.** Summary of transcription factor binding site (TFBS) enrichment and conservation analysis for each module using MEME and *Tomtom*.

**Table S9.** Gene annotation enrichment of each module in 55 main functional categories (P value < 0.05).

**Table S10**. Consensus functional terms associated with GO enrichment and enriched TF binding sites for module target genes.

**Table S11.** Summary of *F. graminearum* and *S. cerevisiae* BLASTp results using predicted top regulators. Only significant hits are listed (E-value < 1e-5).

**Table S12.** Phenotype association analysis for each module.

**Table S13.** Comparison of the virulence TRNs with virulence-associated genes in the PHI-base database

**Table S14.** Regulatory networks compartmentalization analysis using Fisher’s exact test. FS: *F. graminearum*-specific; Core: Conserved genome. Mix: not enriched.

**Table S15.** Comparing the TRNs with previous *F. graminearum* GRNs (Guo *et al*., 2016)

