## Supplementary Figures for "Network Inference from Multi-omic Data Uncovers Dynamic Transcriptional Regulation Modules in Pathogenic Fungus *Fusarium graminearum*"

**Modularized Transcriptional Regulatory Networks of *Fusarium graminearum***

**This PDF file includes:**

Fig. S1-1 to Fig. S1-49 (Figures for all 49 regulatory modules)

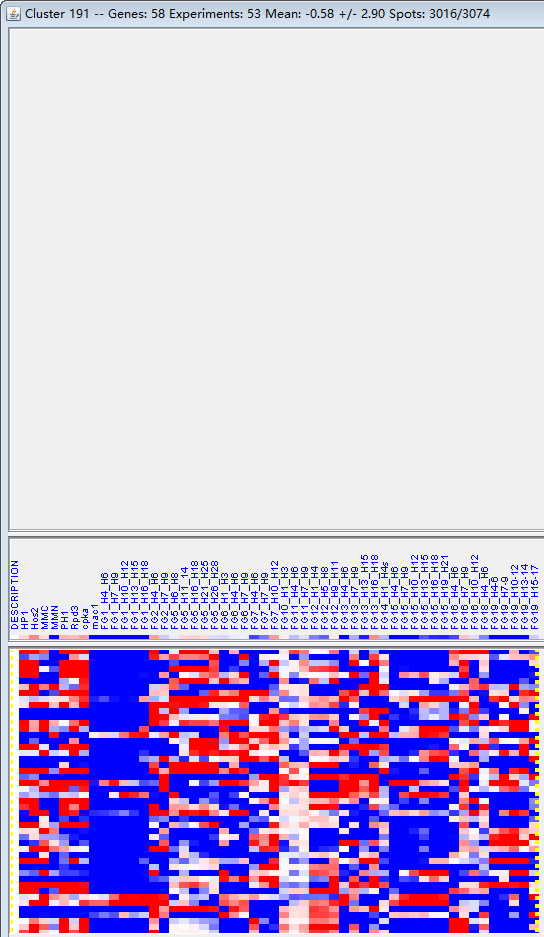

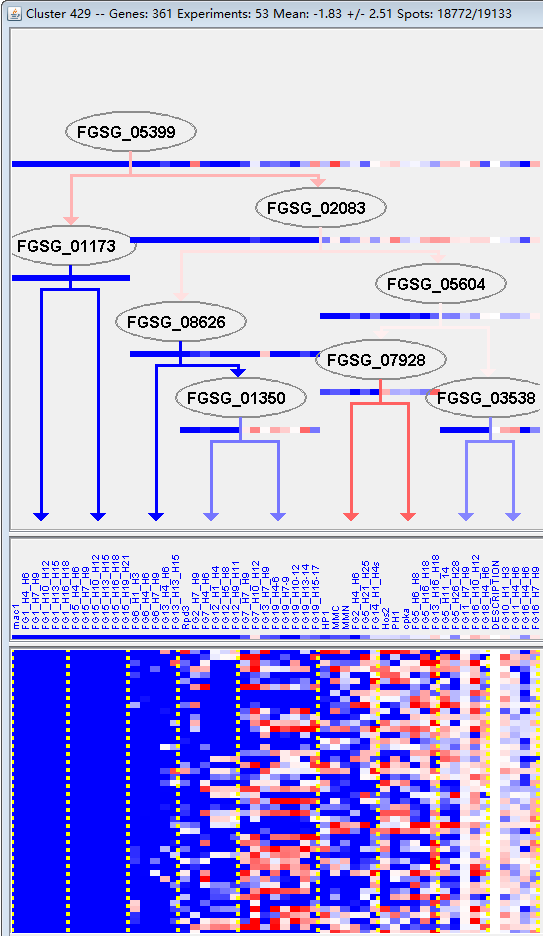

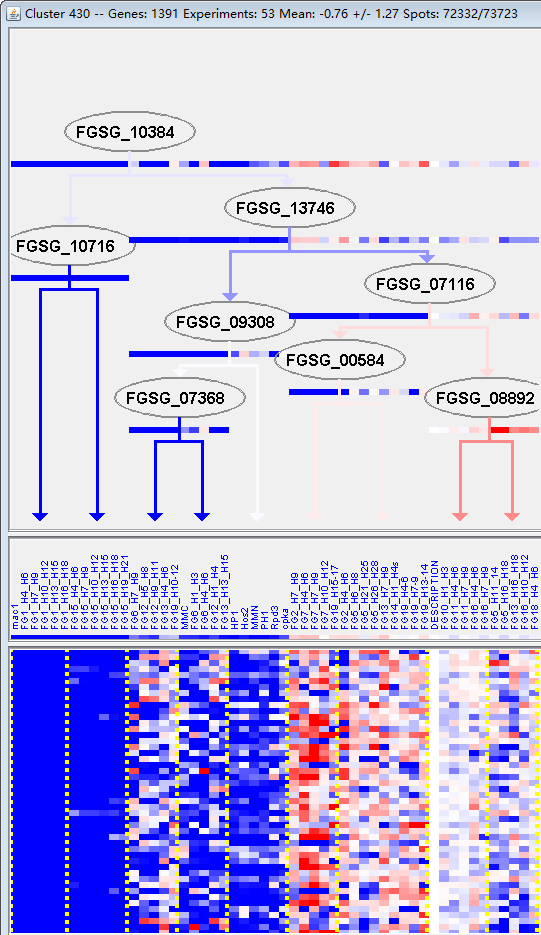

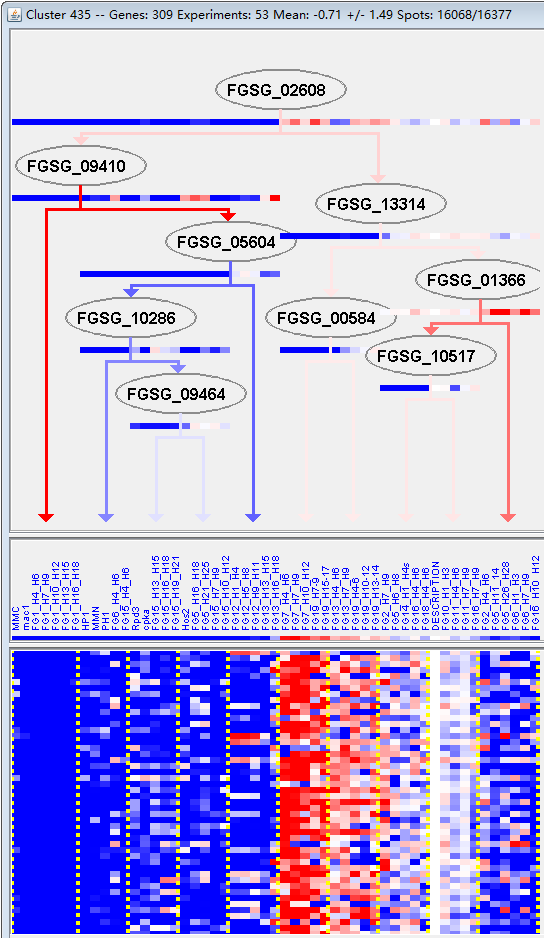

**M1**

**M2**

**M3**

**M4**

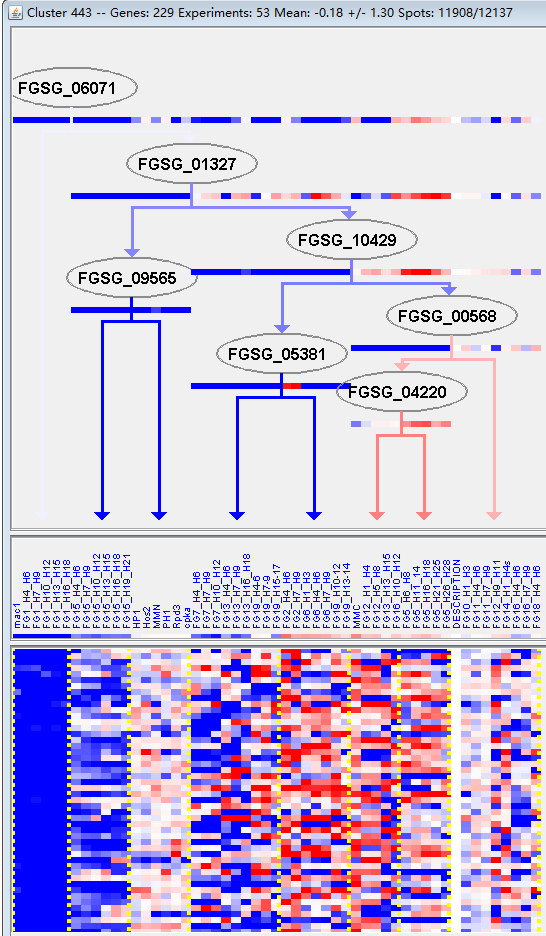

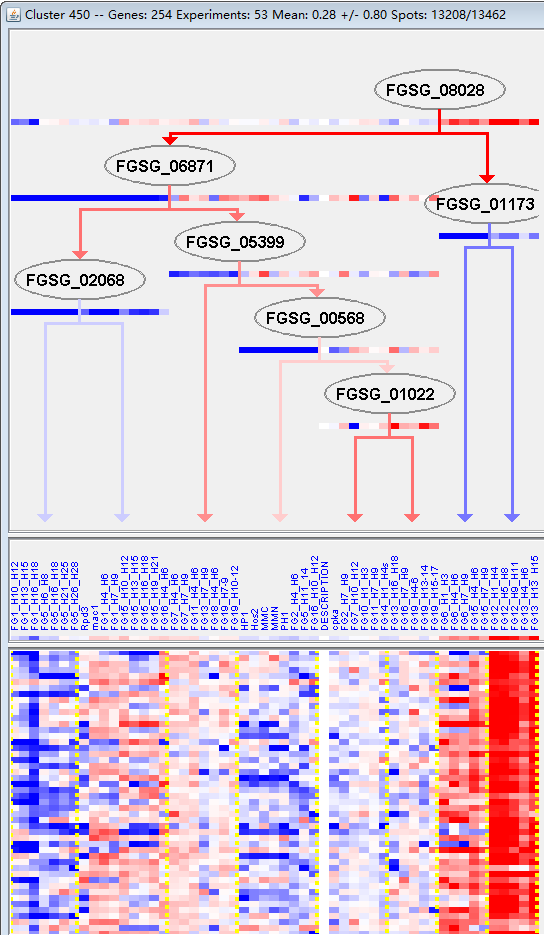

**M5**

**M6**

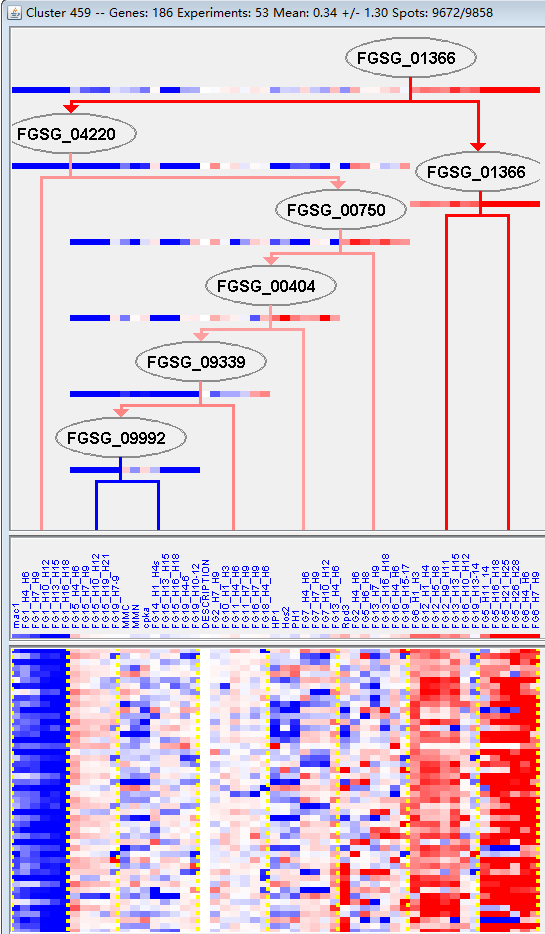

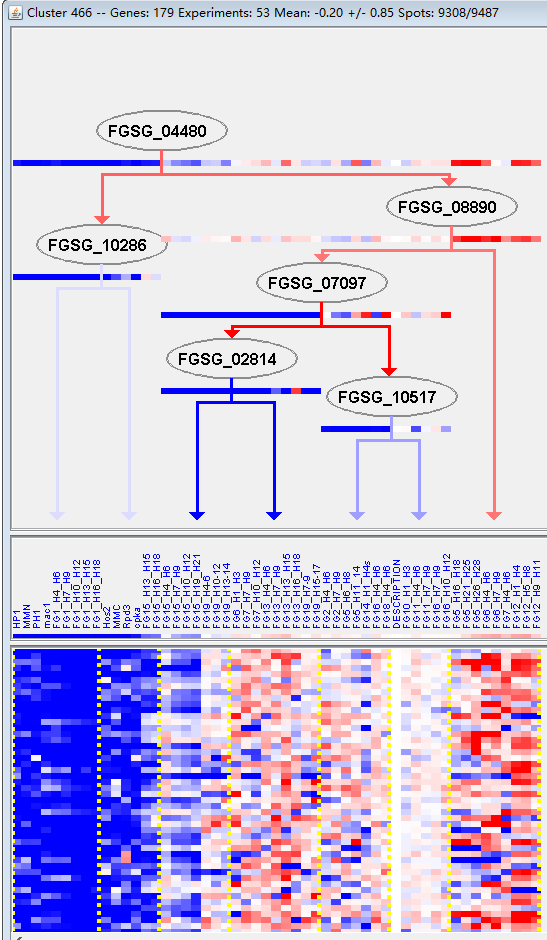

**M7**

**M8**

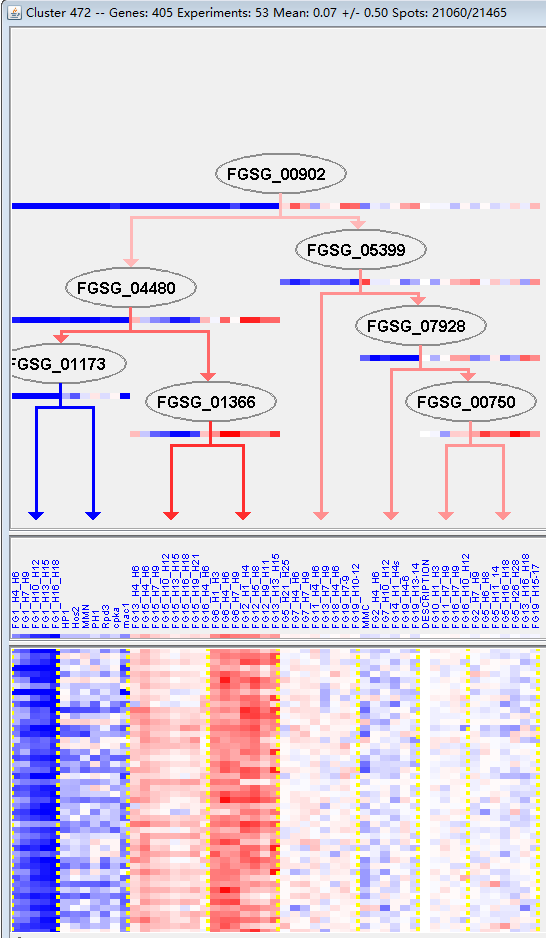

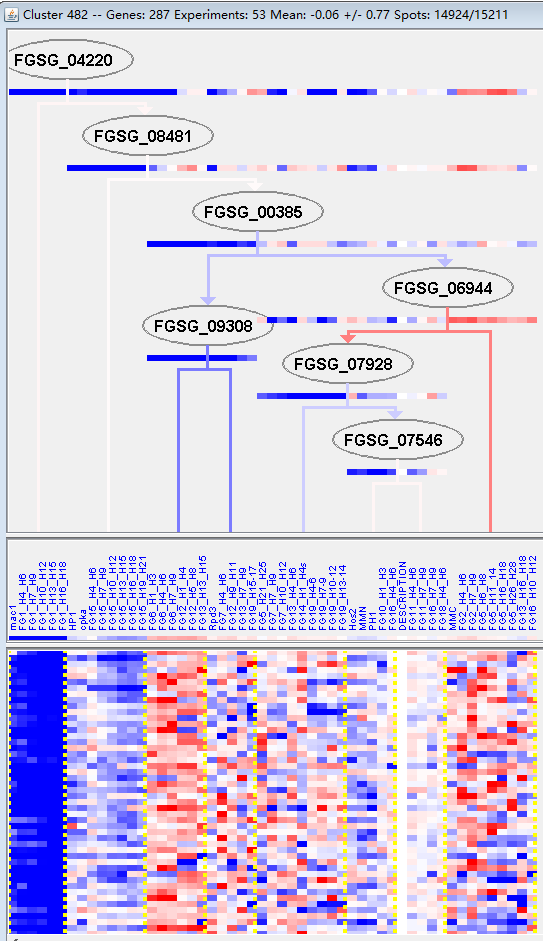

**M9**

**M10**

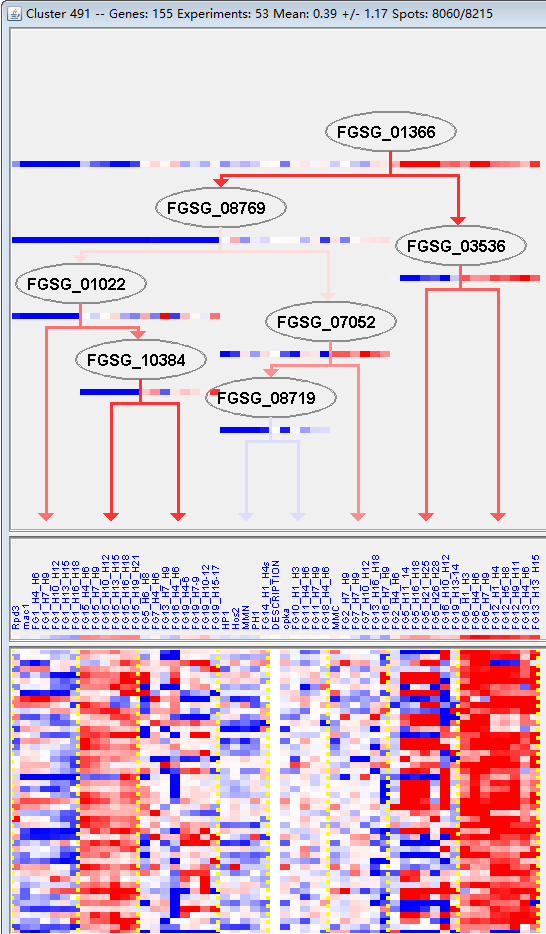

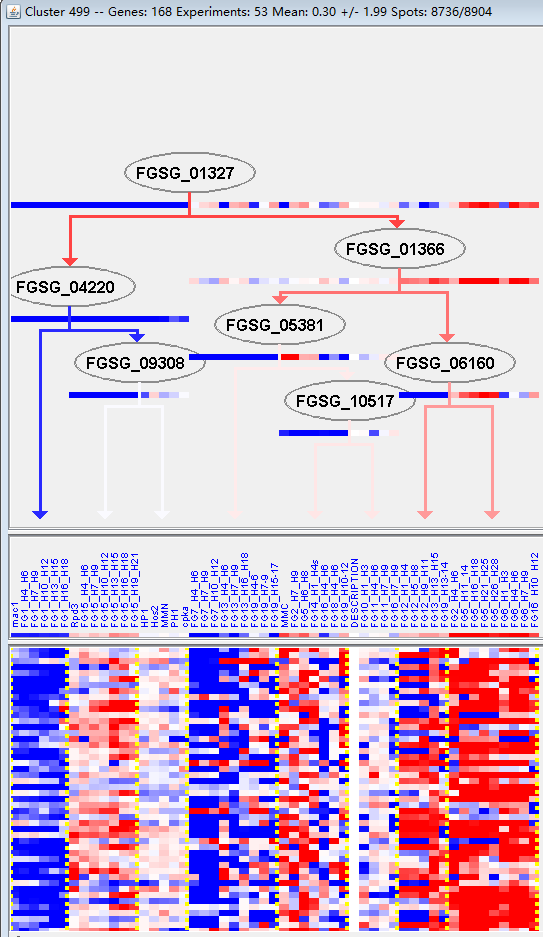

**M11**

**M12**

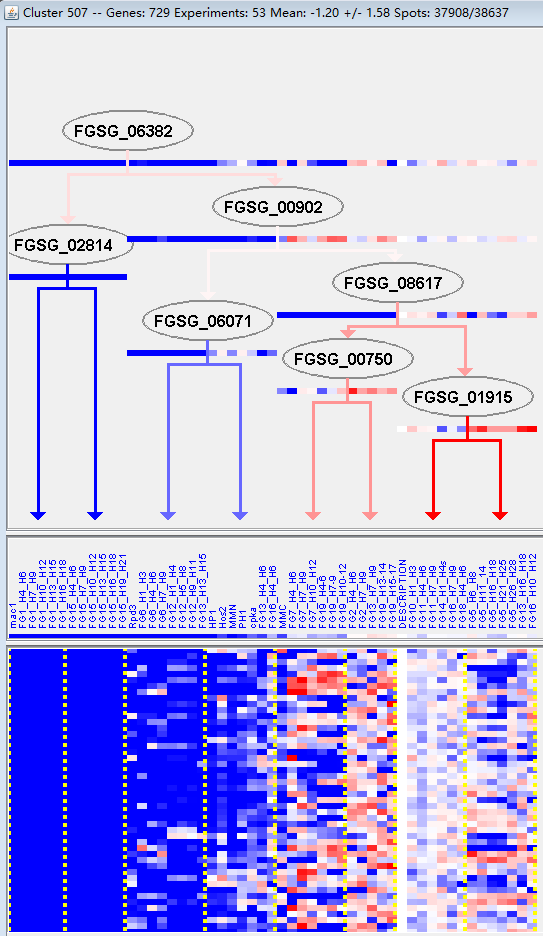

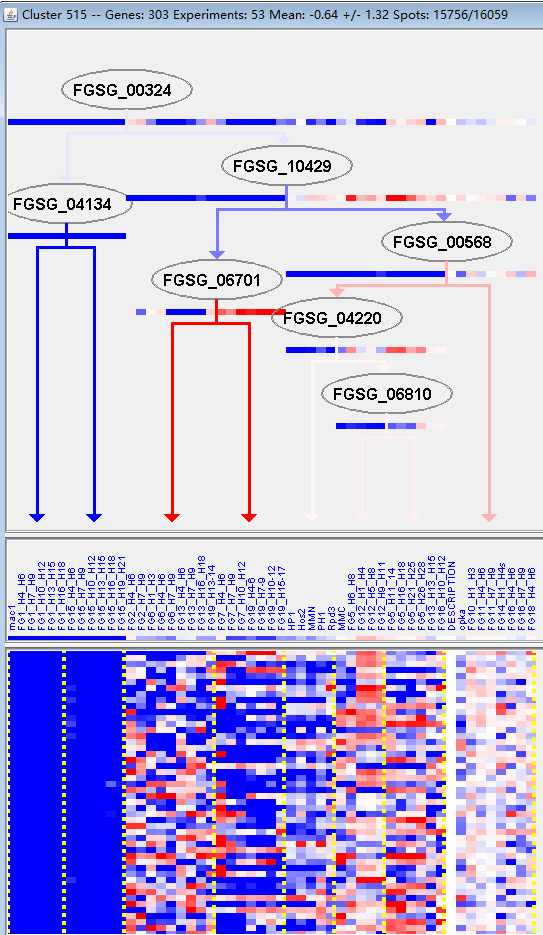

**M13**

**M14**

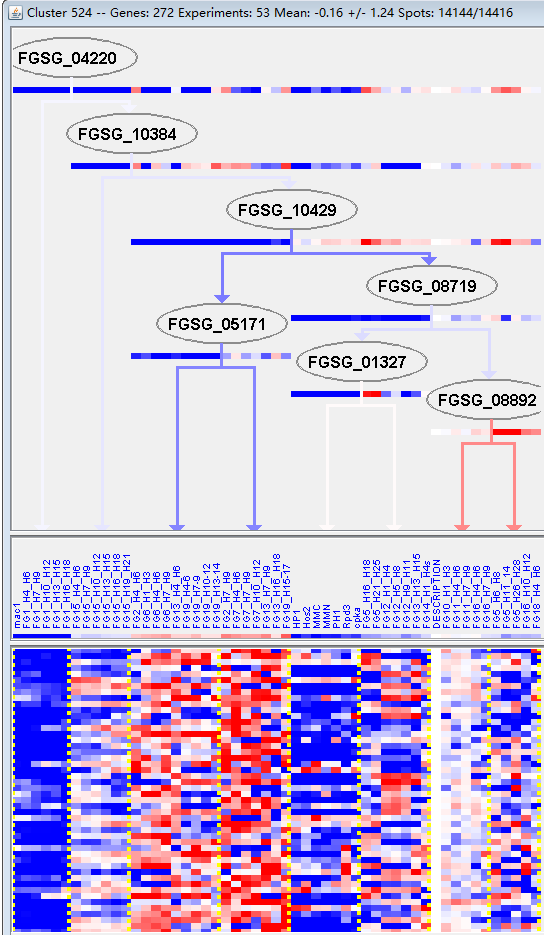

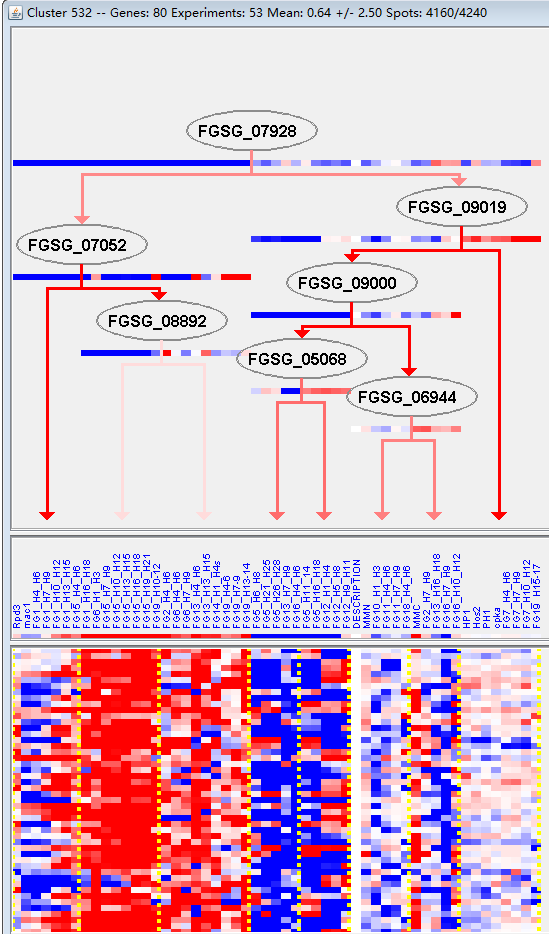

**M15**

**M16**

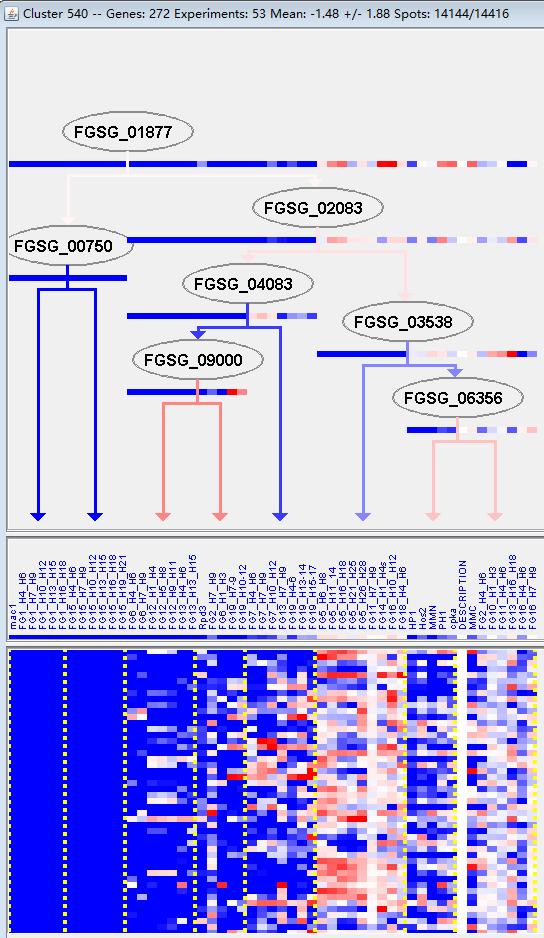

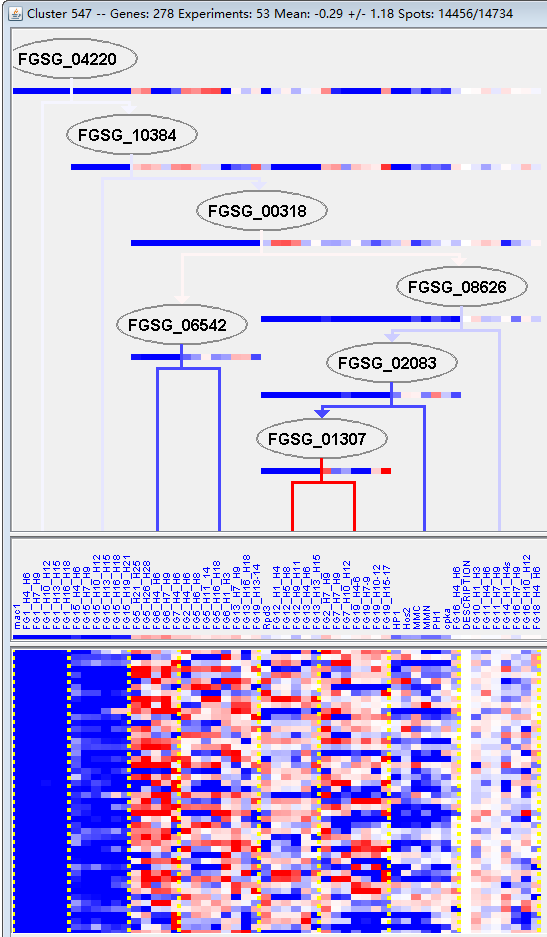

**M17**

**M18**

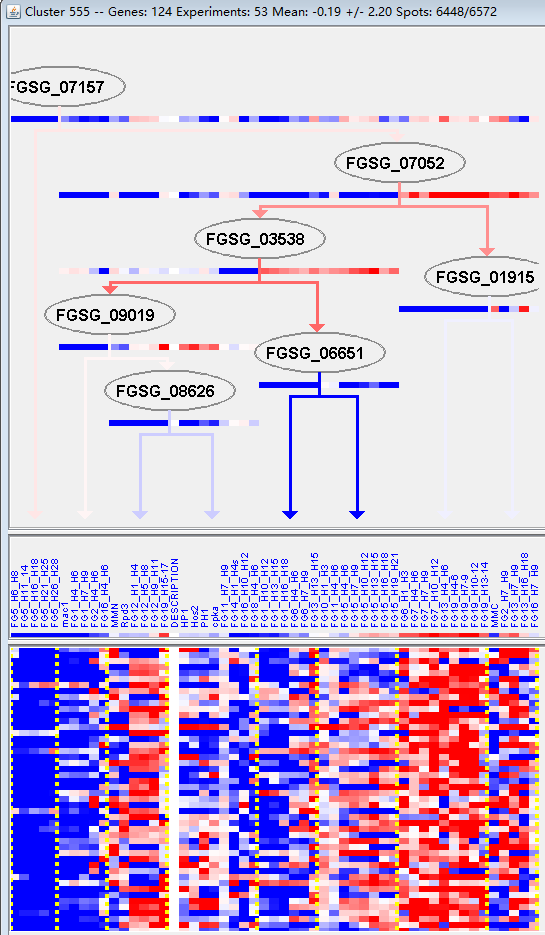

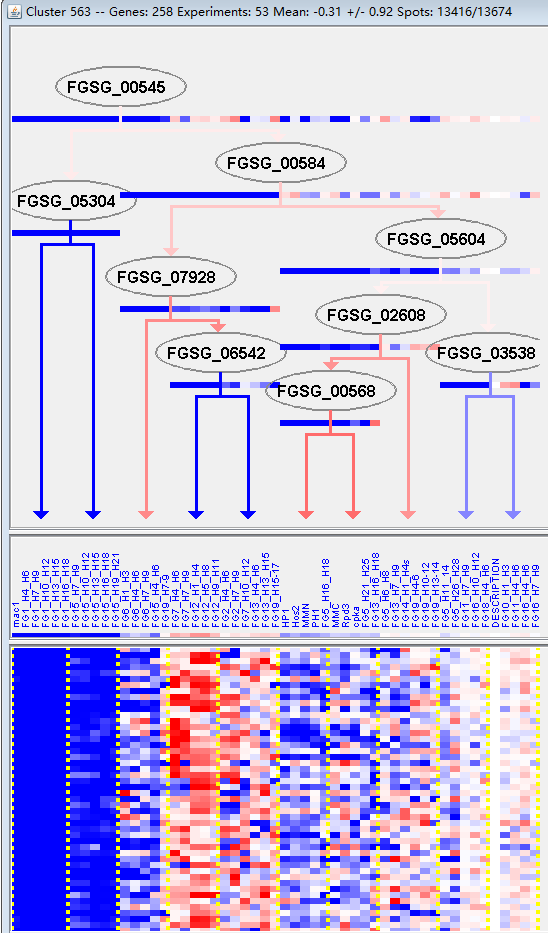

**M19**

**M20**

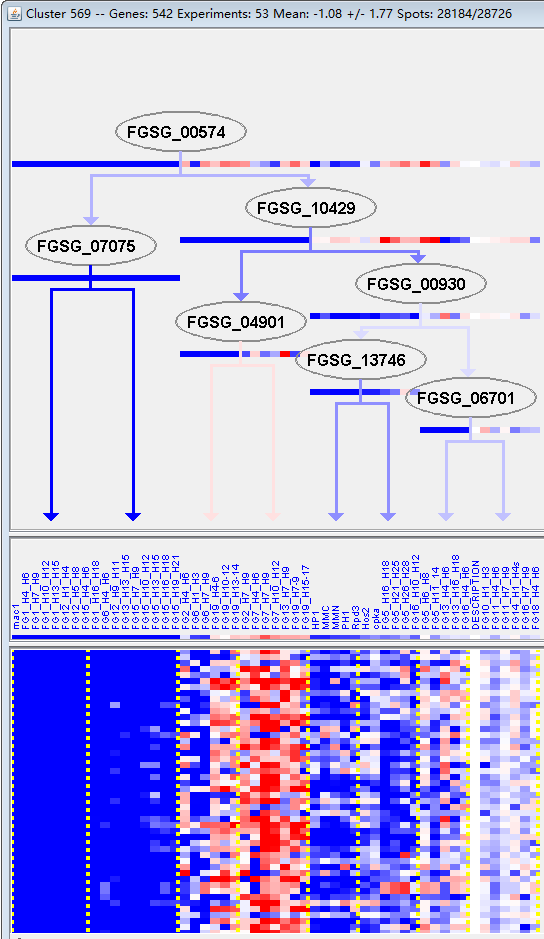

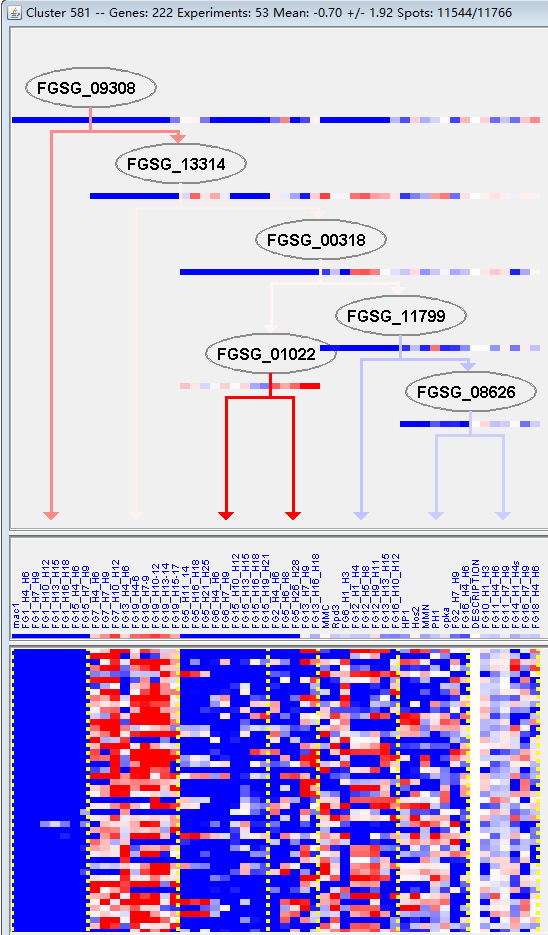

**M21**

**M22**

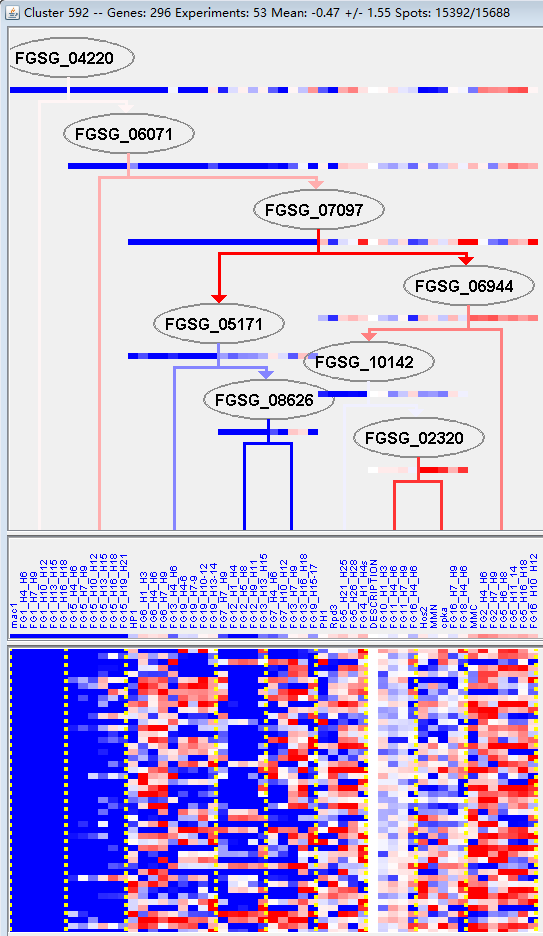

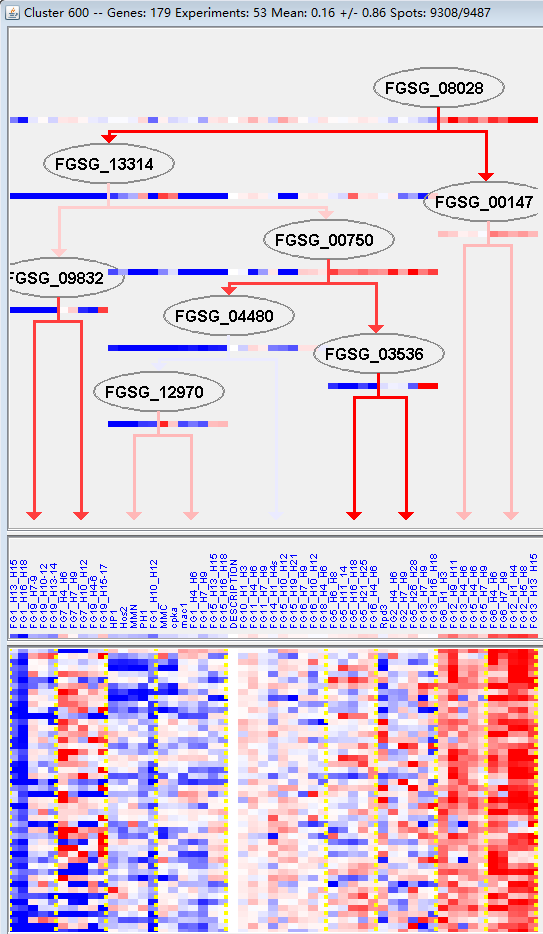

**M23**

**M24**

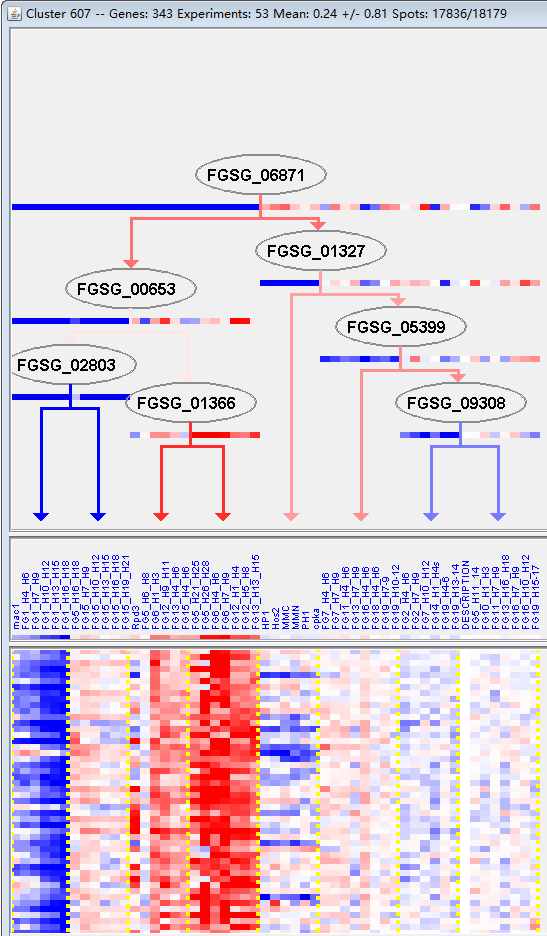

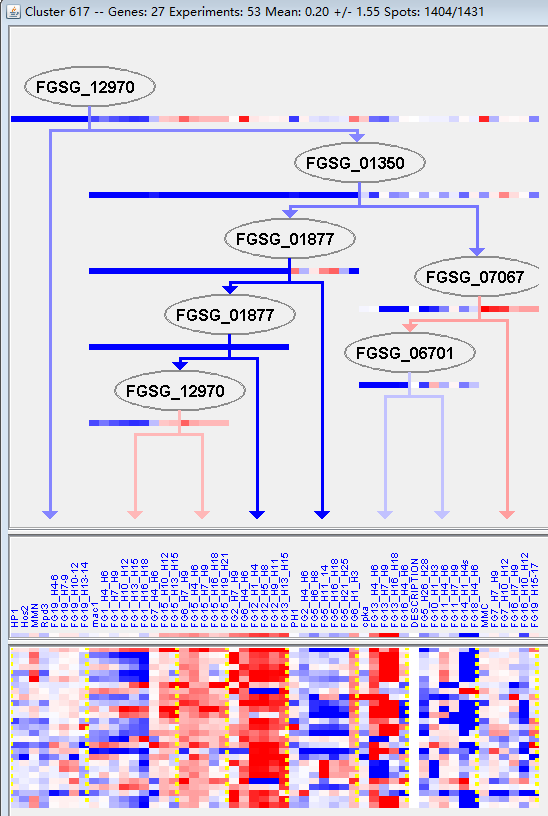

**M25**

**M26**

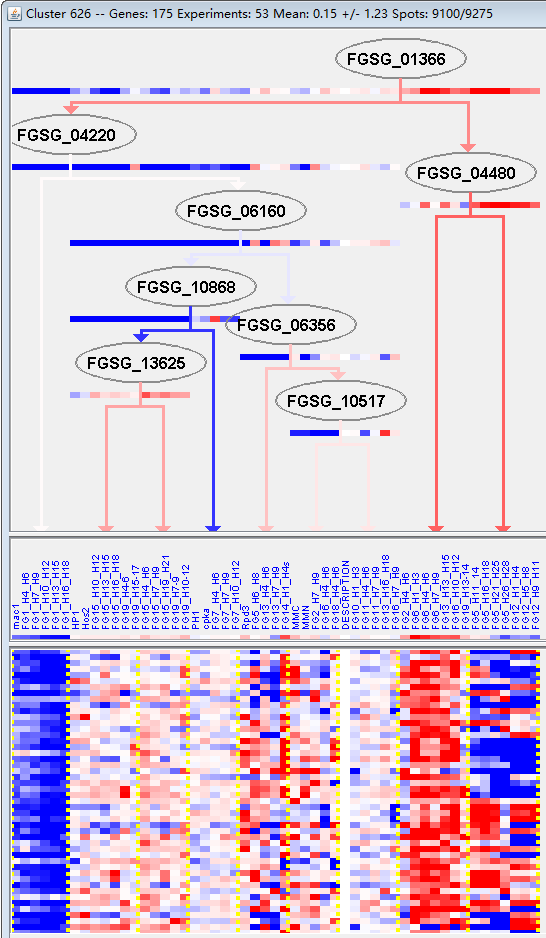

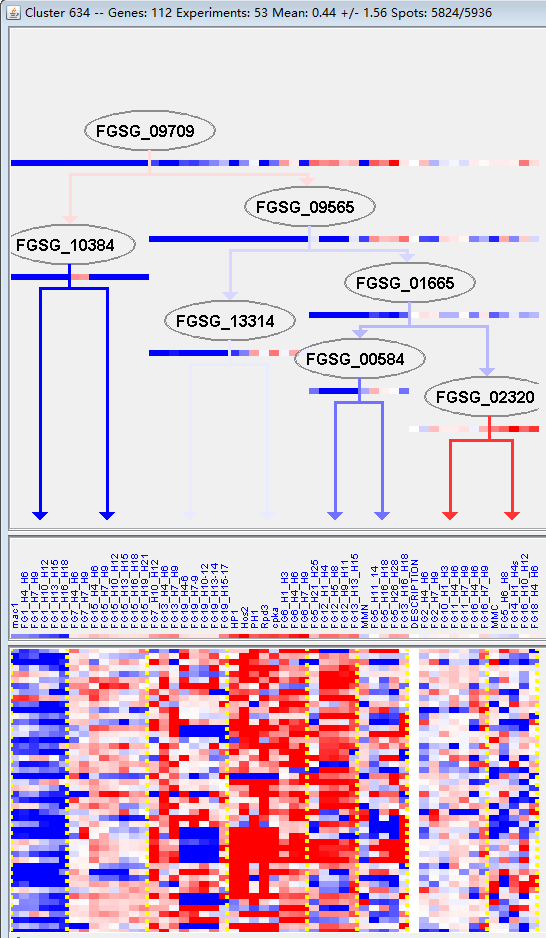

**M27**

**M28**

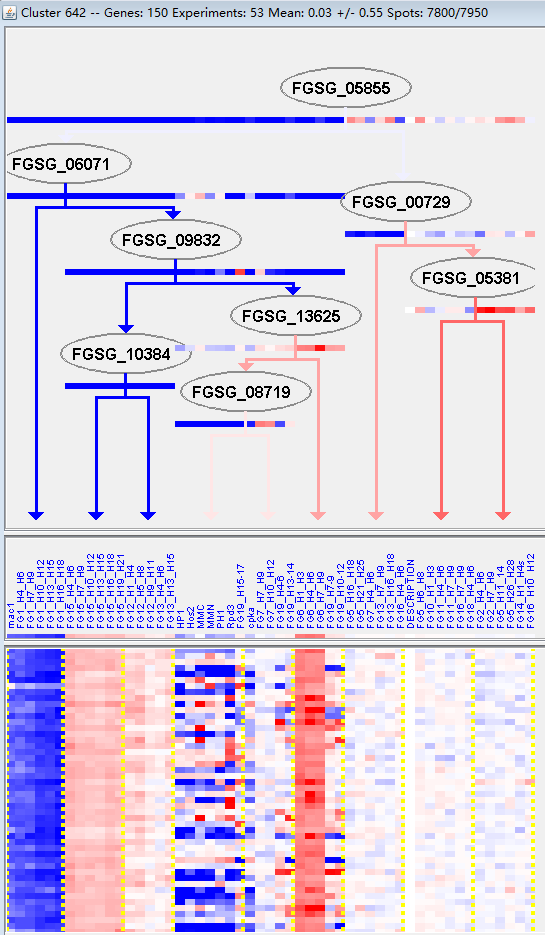

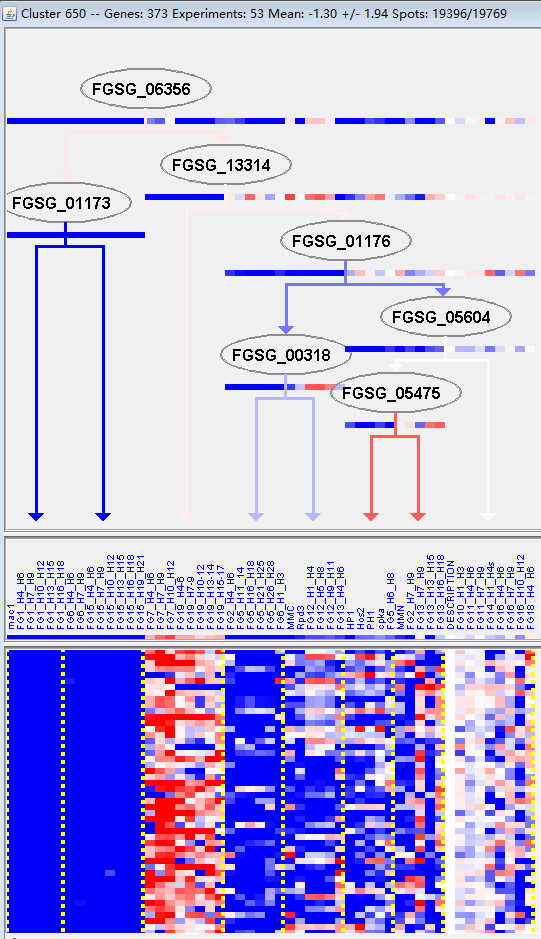

**M29**

**M30**

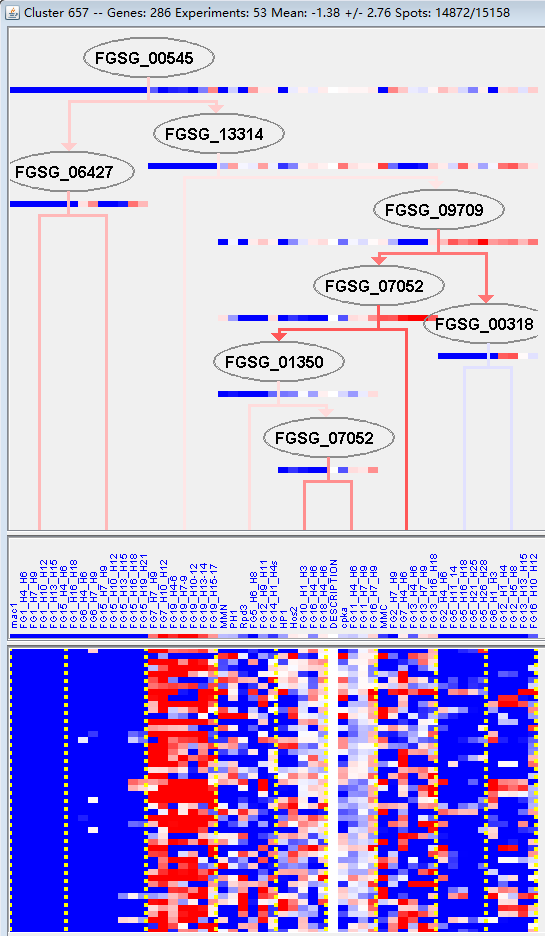

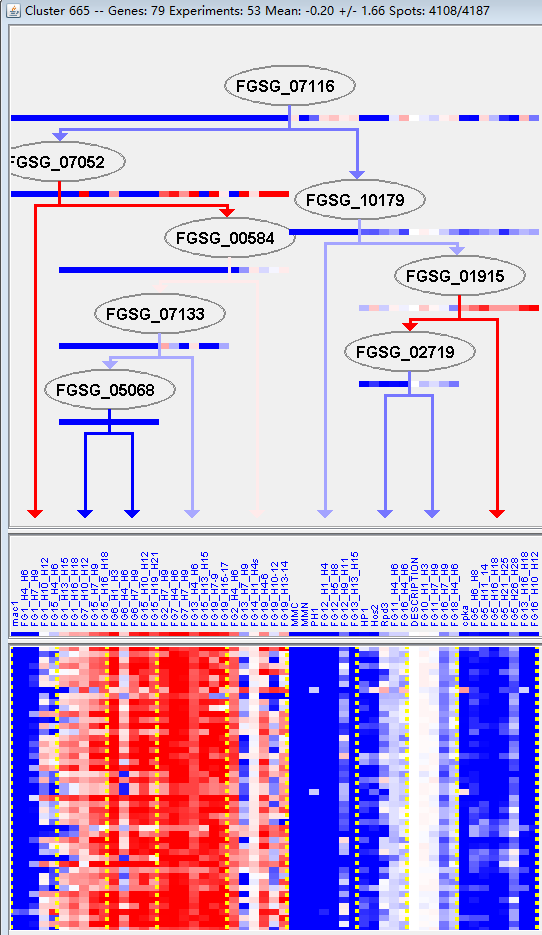

**M31**

**M32**

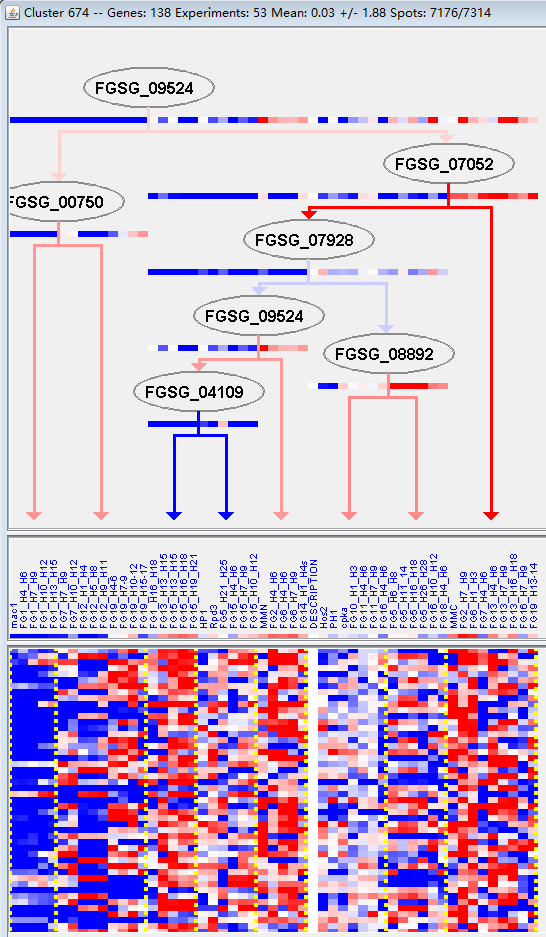

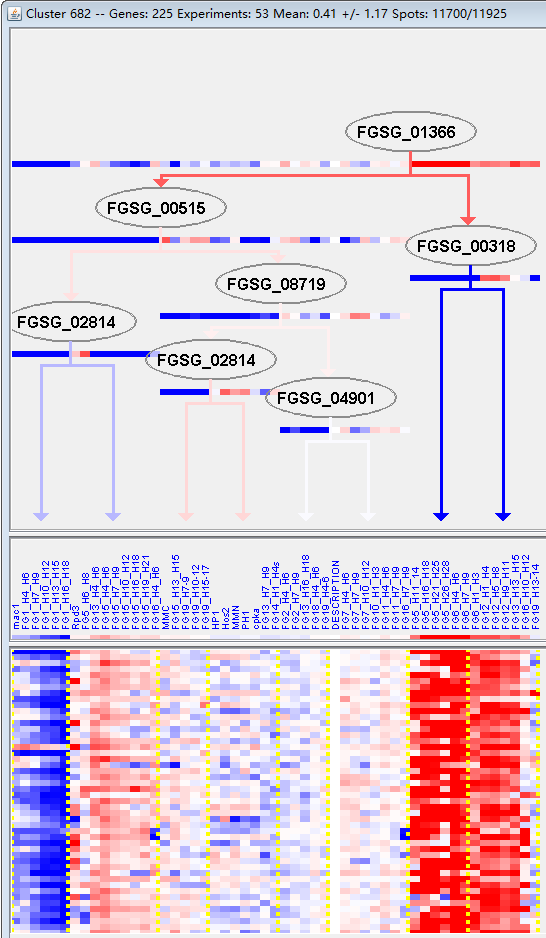

**M33**

**M34**

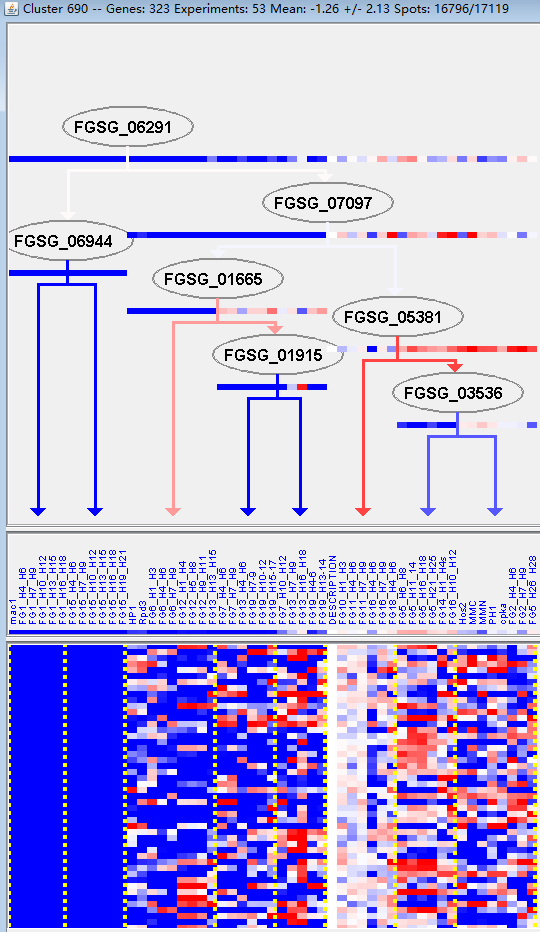

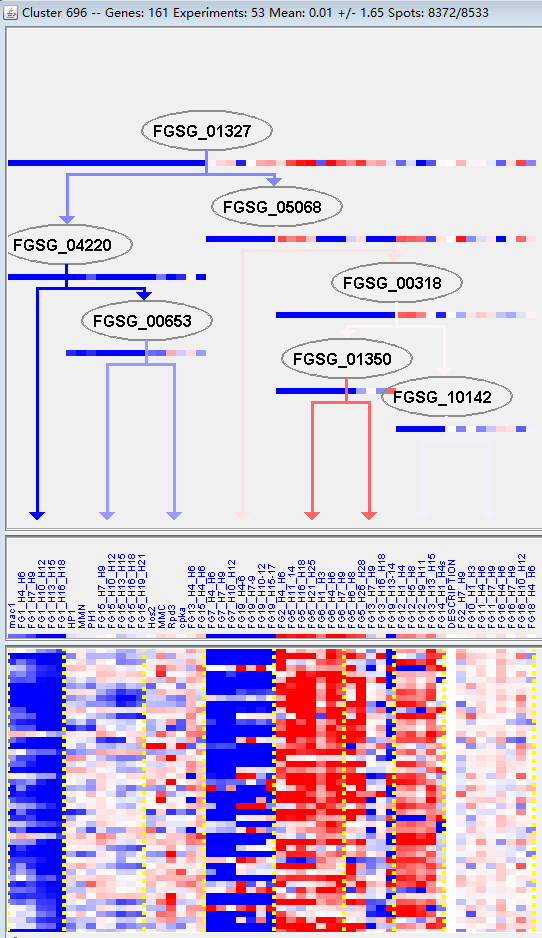

**M35**

**M36**

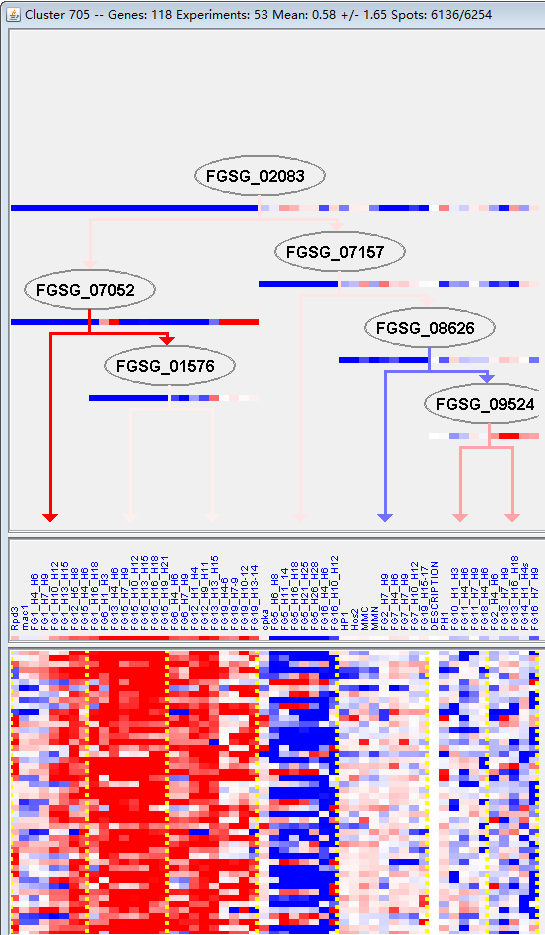

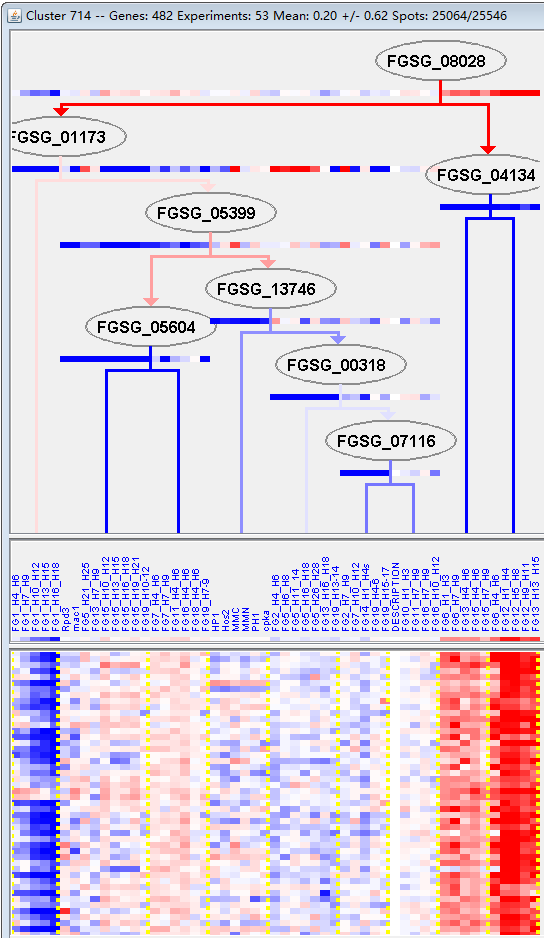

**M37**

**M38**

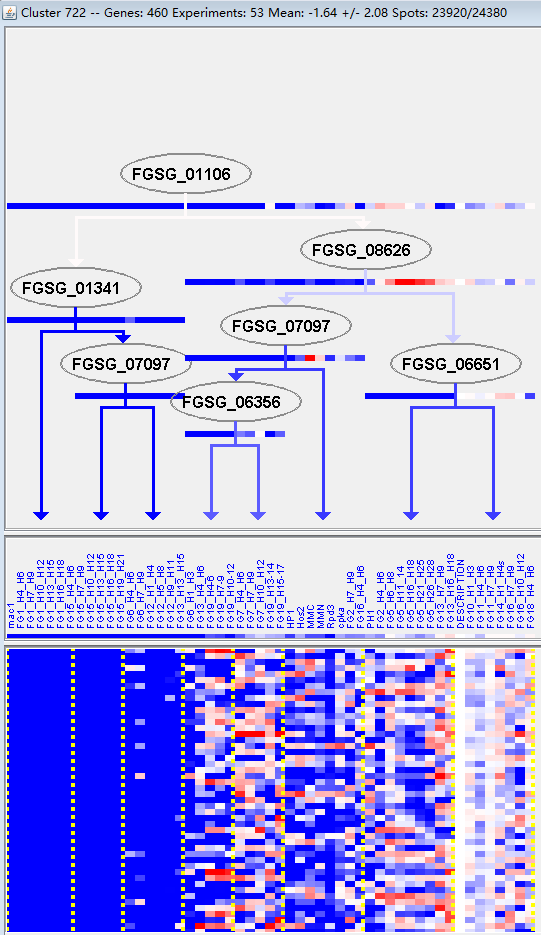

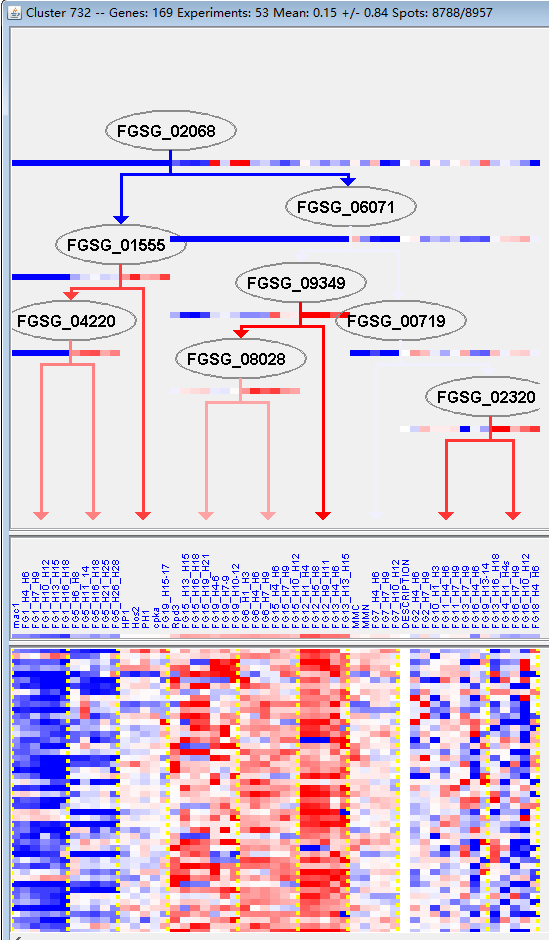

**M39**

**M40**

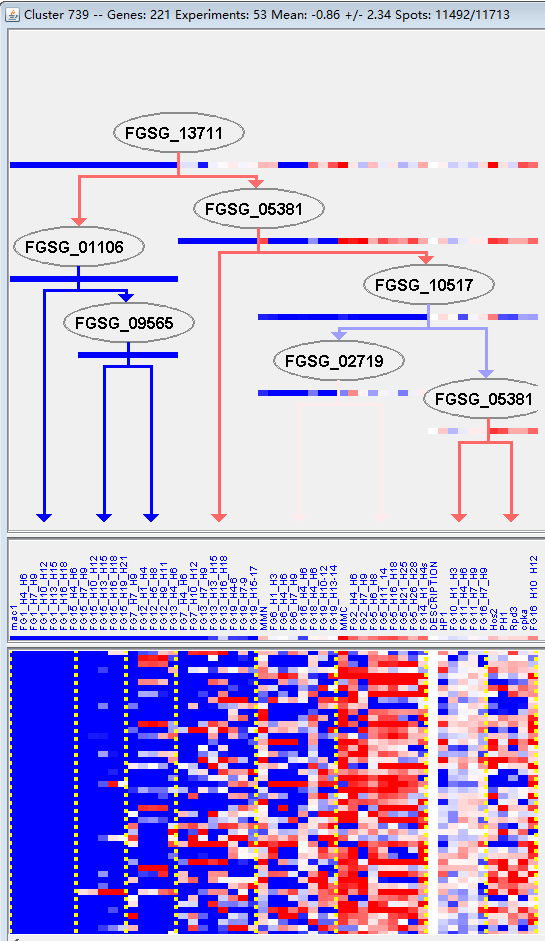

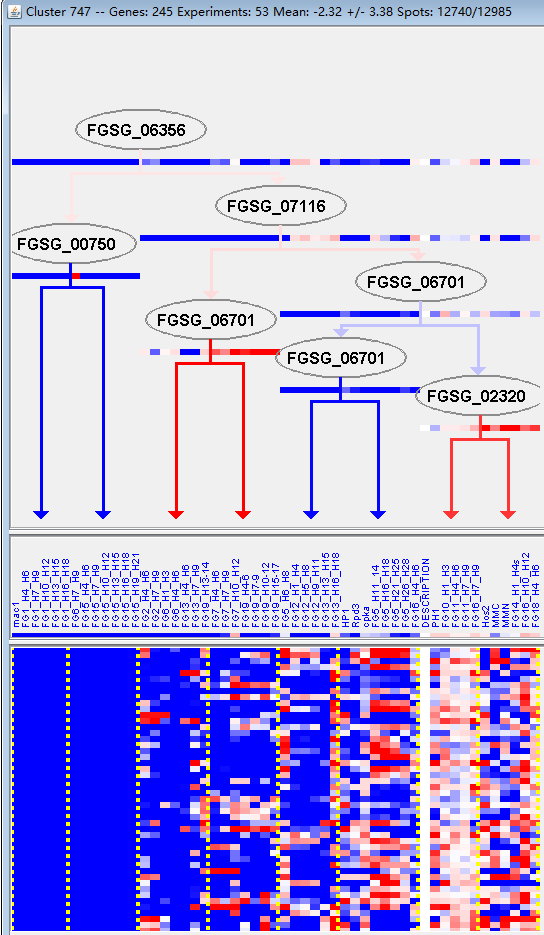

**M41**

**M42**

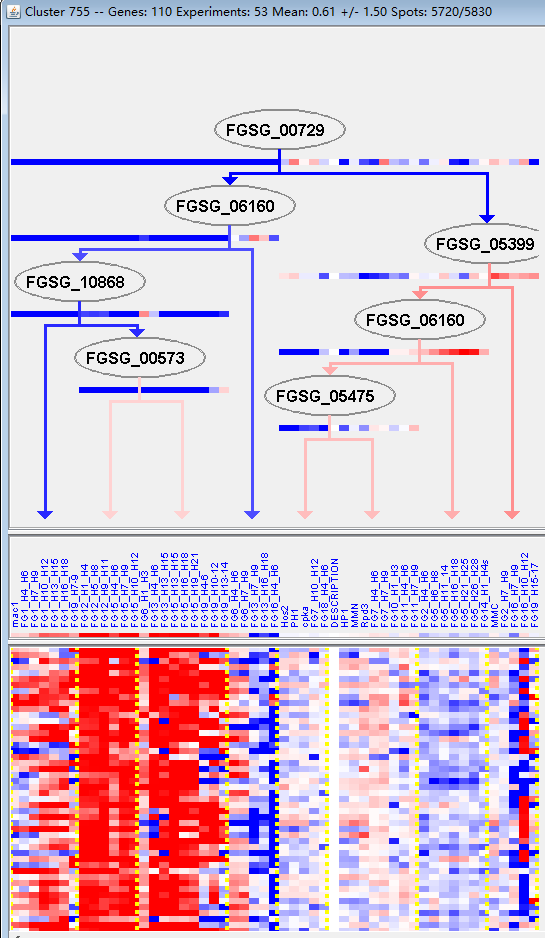

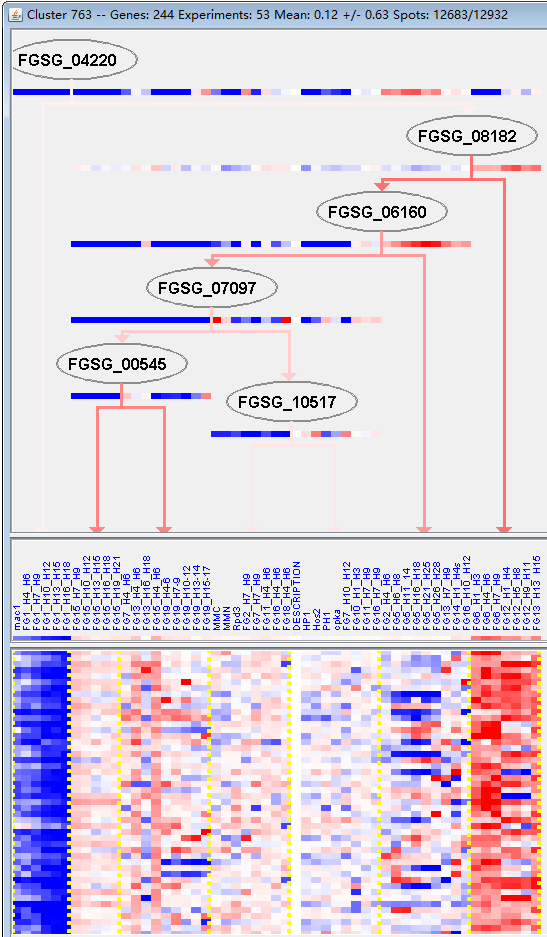

**M43**

**M44**

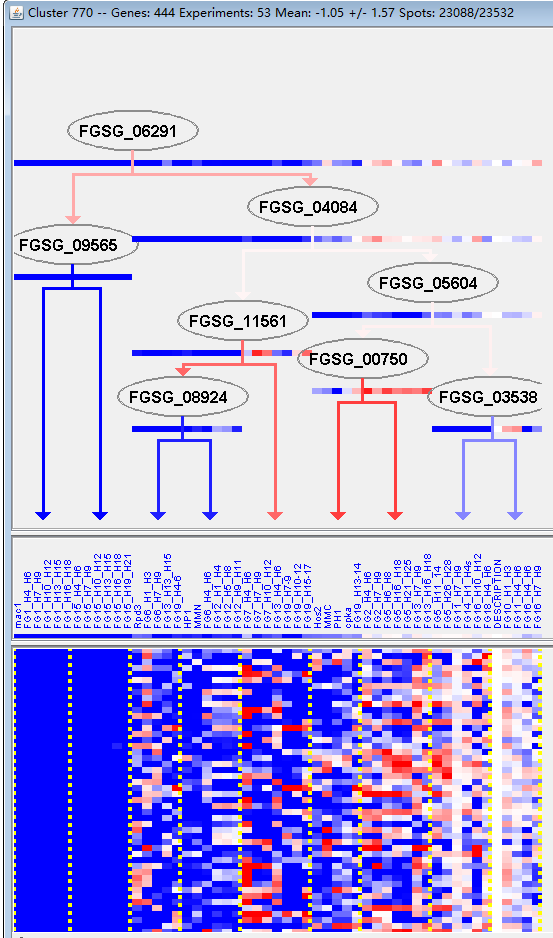

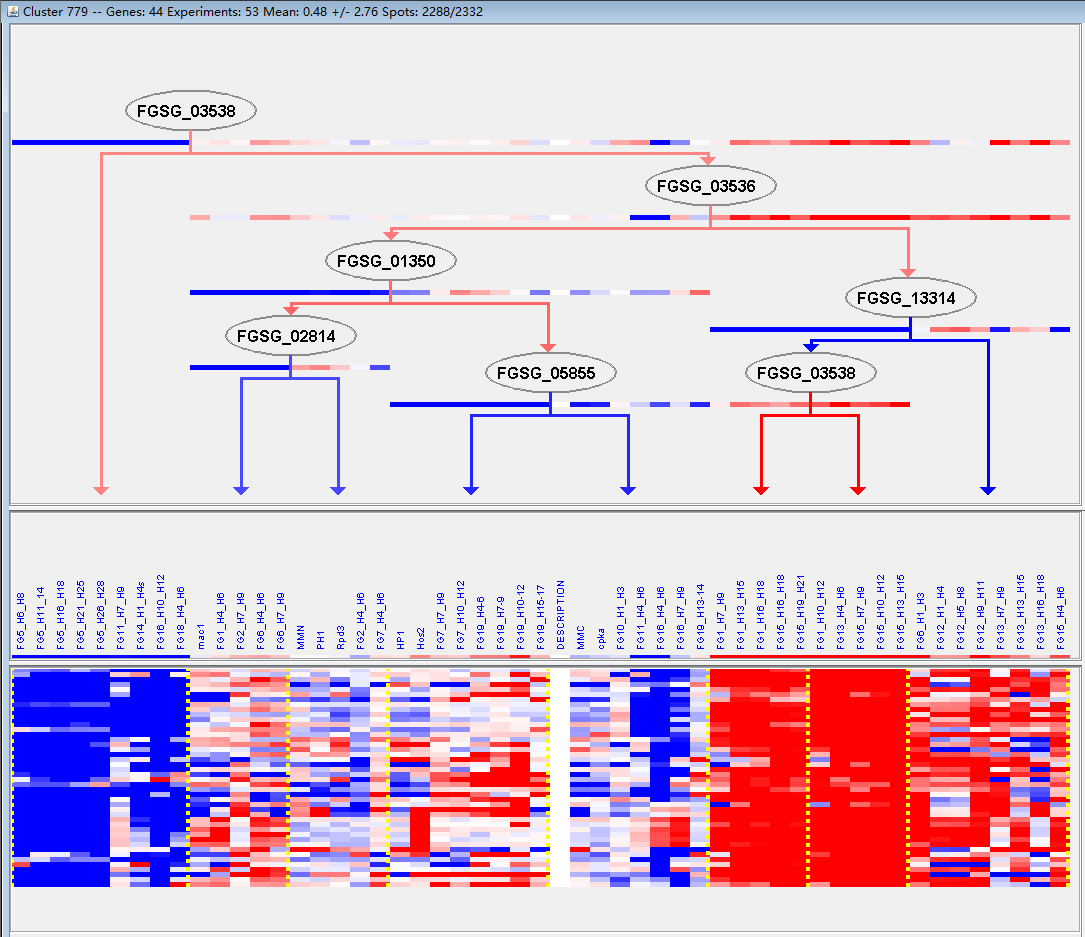

**M45**

**M46**

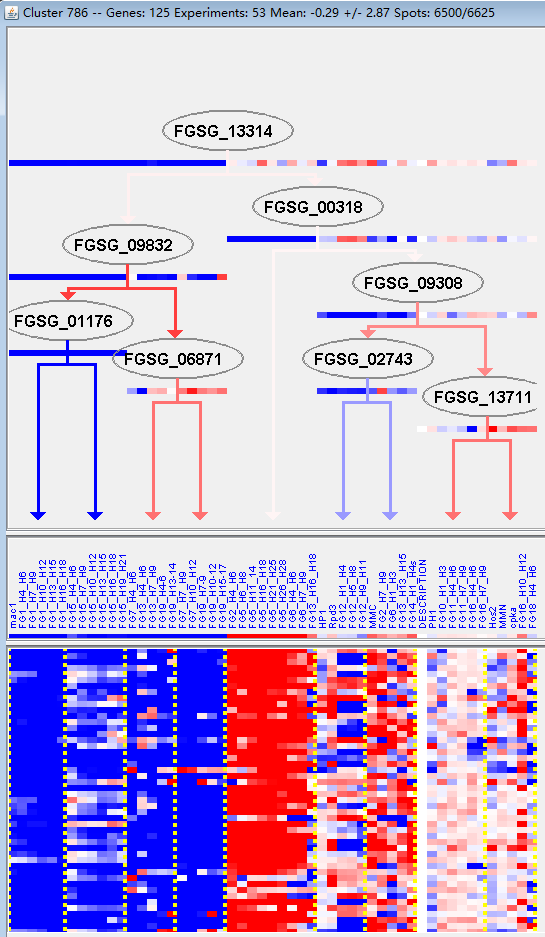

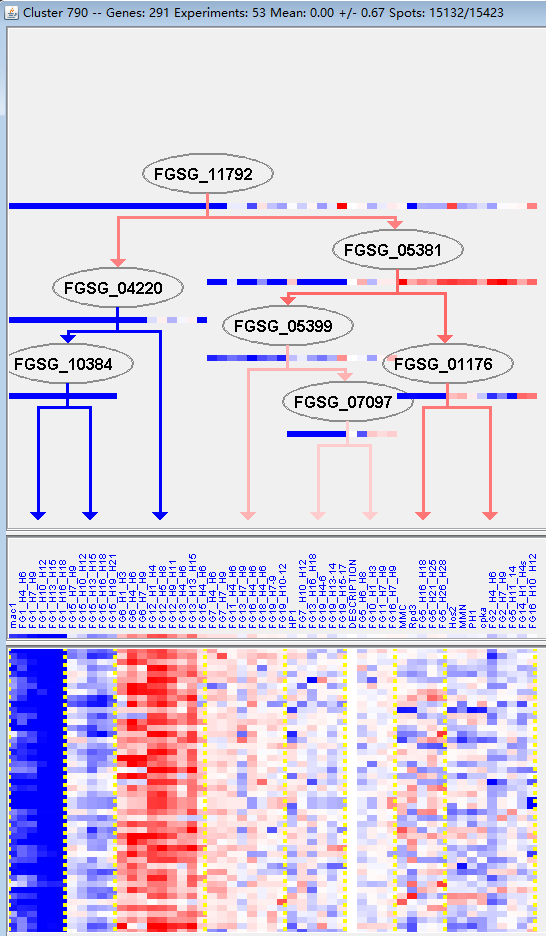

**M47**

**M48**

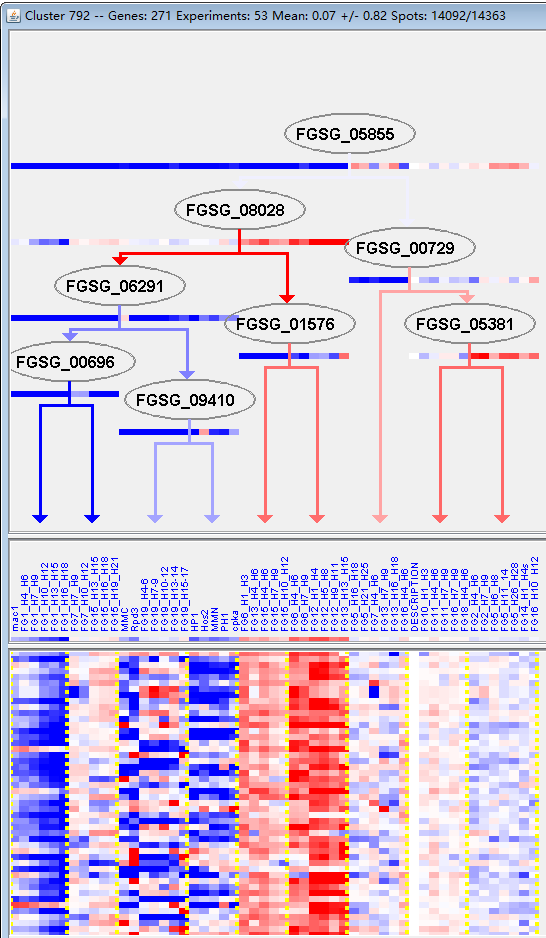

**M49**
